# Contrasting species functional trait structuring of subarctic versus subtropical copepod communities

**DOI:** 10.1101/2020.01.31.928705

**Authors:** Carmen García-Comas, Chih-hao Hsieh, Sanae Chiba, Hiroya Sugisaki, Taketo Hashioka, S. Lan Smith

## Abstract

Classic niche theory assumes that species-level functional traits affect species relative fitness and thus community structuring, but empirical tests of this assumption are scarce. Moreover, recent evidence shows increasing functional over-redundancy towards the tropics, suggesting that the extent to which functional traits confer species’ fitness and thus impact community structuring differs across latitudes. Here, we develop a new method: comparing the frequencies of trait categories in the species-rank abundance distributions of local communities versus their frequencies in the regional average species pool. We contrasted subarctic versus subtropical copepod communities for six important traits. In subarctic communities, medium-sized and cold-water species are selected to dominate, thus traits affect relative fitness as predicted by classic niche theory. In subtropical communities, most species are small and warm-water, but these categories are not selected to dominate, suggesting that greater diversity towards the tropics results from lesser trait-based fitness differences allowing more species to coexist.

## Main

Research on species-level functional traits abounds due to the prevailing concept that species traits play key roles in structuring communities and determining functioning such as predator-prey interactions and trophic transfer^1–3^. Functional traits are defined as the characteristics of an organism that affect its fitness, and subsequently, by increasing the individual ability to grow, survive and reproduce, functional traits affect species abundance (population size) in a community^3^. In general, communities are usually made of few dominant and many rare species^4^. According to classic niche theory, a species is selected to dominate because it possesses functional trait values that favour its niche occupation and confer greater fitness compared to other species in the community ^5,6^. In addition, to explain the existence of many rare species, classic niche theory predicts species coexistence by niche segregation: species avoid competition by avoiding niche overlap ^5,7^. However, modern niche theory adds that species coexistence can still occur among species of overlapping niches (i.e., with similar functional trait values) if these species have similar fitness ^8^. Finally, disregarding trait effects on community structuring, neutral theory argues that species dominance occurs mainly due to stochastic processes such as drift and dispersal^9^. Which gives rise to an important question that we intend to assess: Can species traits explain species dominance and coexistence in natural communities?

A myriad of trait-based indices have been proposed e.g., ^2,12,13–15^, including various facets of diversity (i.e., richness, evenness and divergence) ^15,16^, albeit that many of them are redundant ^17^. The most popular indices are community-integrated measures such as the community weighted mean (CWM), Convex Hull Volume ^13^ and Rao quadratic entropy ^12^. These indices have proven useful to characterise how community traits change responding to environmental conditions e.g., ^18,19^ and affect functioning e.g., ^20^. Yet, community-integrated trait indices may not be suitable to answer whether and how traits structure species communities. Community-integrated indices assume that all of the selected traits have the same weight in community structuring or functioning. However, functional traits should not be considered similar in structuring communities; instead, we should select and weight traits to better define niches ^1^. To determine the important functional traits for community structuring, Maxent and Traitspace models, developed to predict species abundances from environment-trait relationships, have the potential to detect important traits to define species niches; however, the selection criteria are based on processes of environmental filtering but neglect species interactions ^21^. Moreover, these models predict species relative abundances, according to the species trait values, by statistically fitting the CWM for those traits, and thus they rely on a community-integrated index.

Community-integrated trait indices do not inform whether the observed variation in trait composition results from changes in the proportion of species with certain trait values (i.e., relative richness) or from changes in the abundance of species (i.e., relative abundance). This issue is critical if we aim to understand the role of traits in community structuring (i.e., species relative fitness), or how the species coexist according to their trait composition. Indeed, recent global biogeographical studies of mammals and fishes, applying community-integrated indices of species-level functional traits (i.e., for each trait, assigning a single trait value to all individuals belonging to a species) report uncoupling of functional and species diversity, with apparent functional redundancy towards the tropics ^22–24^. But the *integrated* nature of these indices prevented detailed investigation of how traits affect community structuring.

We hypothesize that the role of functional traits on species community structuring varies among traits and across latitudes. Specifically, elevated functional redundancy towards the tropics points to denser species niche packing towards those regions. Thus, following modern niche theory, we hypothesize that the effect of functional traits on species fitness is dampened towards the tropics, favouring many species with similar niches to coexist ^8^. In order to test if species-level categorical traits (i.e., for each trait, assigning a single category to all individuals belonging to a species) structure communities differently across latitudes, we develop a new method: We add trait information to species-rank abundance distributions (RADs). RAD provides information about species dominance within a community. Our approach provides visual information about the repartition of trait categories along species ranks, allowing us to further interpret patterns and propose hypotheses that cannot be as easily inferred from a null-model approach that lacks this visual information^25^. With our new method, we specifically aim to address a simple, but hitherto neglected, question: Do species-level categorical traits affect the abundance ranking of species and therefore the relative species fitness in a set of communities as expected by classic niche theory? Answering this question allows us to detect if an important trait category is selected to dominate in a region and to compare trait-based community structuring across latitudes; thus, we test the hypothesis-functional traits effects on species fitness dampen towards the tropics.

We apply our method to compare the role of six important traits (body size, diet, feeding mode, myelination, spawning strategy and thermal range; Box 1) in a set of subarctic communities versus a set of subtropical communities of copepods in the western North Pacific. Our method reveals contrasting roles of different traits in structuring subarctic versus subtropical communities. The results partially support our hypothesis that functional traits effects on species fitness dampen towards the tropics. In the broader scope of trait-based ecology, the proposed method could be applied to other systems to unveil generalities that may advance our understanding of community structuring.

## Results and discussion

We apply our method to natural communities of marine copepods. Copepods, as the major group of oceanic mesozooplankton, are a key trophic link between unicellular plankton and fish. However, the first global biogeography of CWM of selected copepod traits has only recently been published ^26^. We considered five important functional traits and one important biogeographic trait for copepods (see their description and importance in Box 1). The six traits are: body size as meta-trait; diet, feeding mode and myelination as trophic traits; spawning strategy as secondary-production trait and thermal range as biogeographic trait. We ought to stress that thermal range is not determined from our dataset but from independent bibliography. Furthermore, we do not consider thermal range as a functional trait: we interpret thermal range as an outcome of several environmentally responsive functional traits that may confer a taxon the specialized ability to inhabit certain environments.

We aim to test if species-level categorical traits affect the abundance ranking (an index of dominancy) of species and therefore the relative species fitness (selection) in a set of subarctic (transported southward within the Oyashio subarctic current) versus a set of subtropical communities (transported northward within the Kuroshio subtropical current). For brevity, hereafter we refer to each set of communities as a metacommunity, although the two sets are not strictly metacommunities because sampling was spatio-temporally heterogeneous throughout 40 years (appendix S1). Specifically, in each metacommunity, we compare the relative frequencies of each trait category in each rank (*F-rank*) with the reference for null selection (*F-pool*) (expectance of relative frequency by chance): the mean relative frequency of species belonging to each category in a local community (i.e., mean relative richness of categories) (see conceptual model in Figure 1 and detailed steps of the method in Figure 2). We interpret *F-rank* as representing current selection: greater frequencies towards the first ranks indicate that the category confers greater fitness (dominance) in a set of communities, because fitness, as the integrated ability to grow, survive and reproduce, is a primary determinant of the relative abundance of species^3^. We interpret *F-pool* (i.e., mean relative richness of categories) as representing legacy effects such as integrated effects of immigration, past selection, adaptation and speciation: accumulation of species belonging to a certain category in a community. In addition, *F-pool* is *per se* a proxy for species niche partitioning with respect to that trait (assuming that the trait affects species niche).

**Figure 1.**
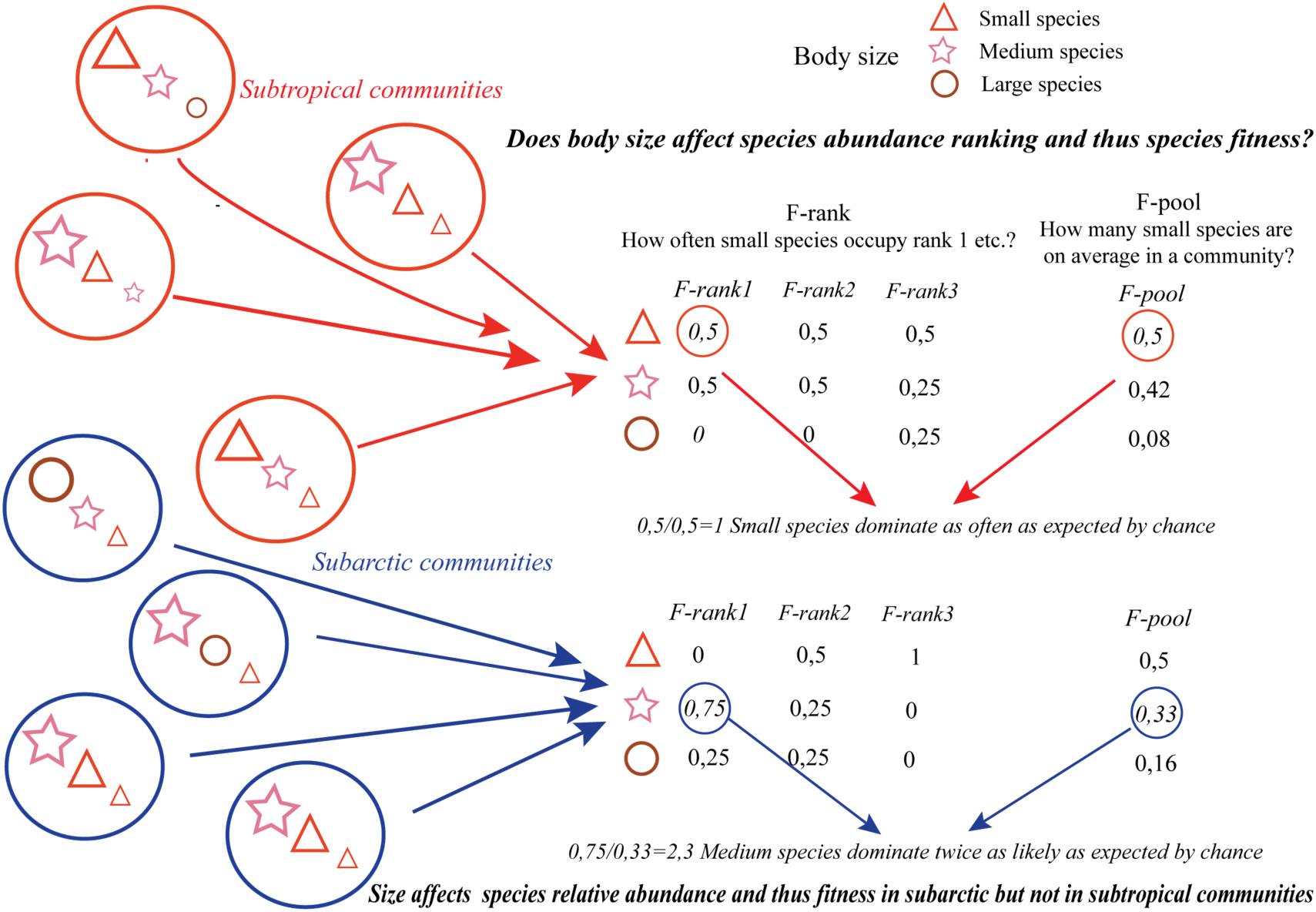
Conceptual diagram of the method: We estimate how often a trait category occupies rank 1 in a set of communities (*F-rank_1_*: relative dominance) (the biggest symbol inside a circle corresponds to the most abundant category in a community). Then, test if this frequency is greater than the average relative presence of species with that trait category in a community (*F-pool*: relative richness) (i.e., we test if the category occupies rank 1 more often than it is represented in the community: more often than expected by chance). If the frequency of occupying rank 1 is greater than the frequency of species with that trait category (*F-rank_1_*/*F-pool*>1), we deem that the trait category confers greater fitness and thus greater relative abundance to a species.

**Figure 2.**
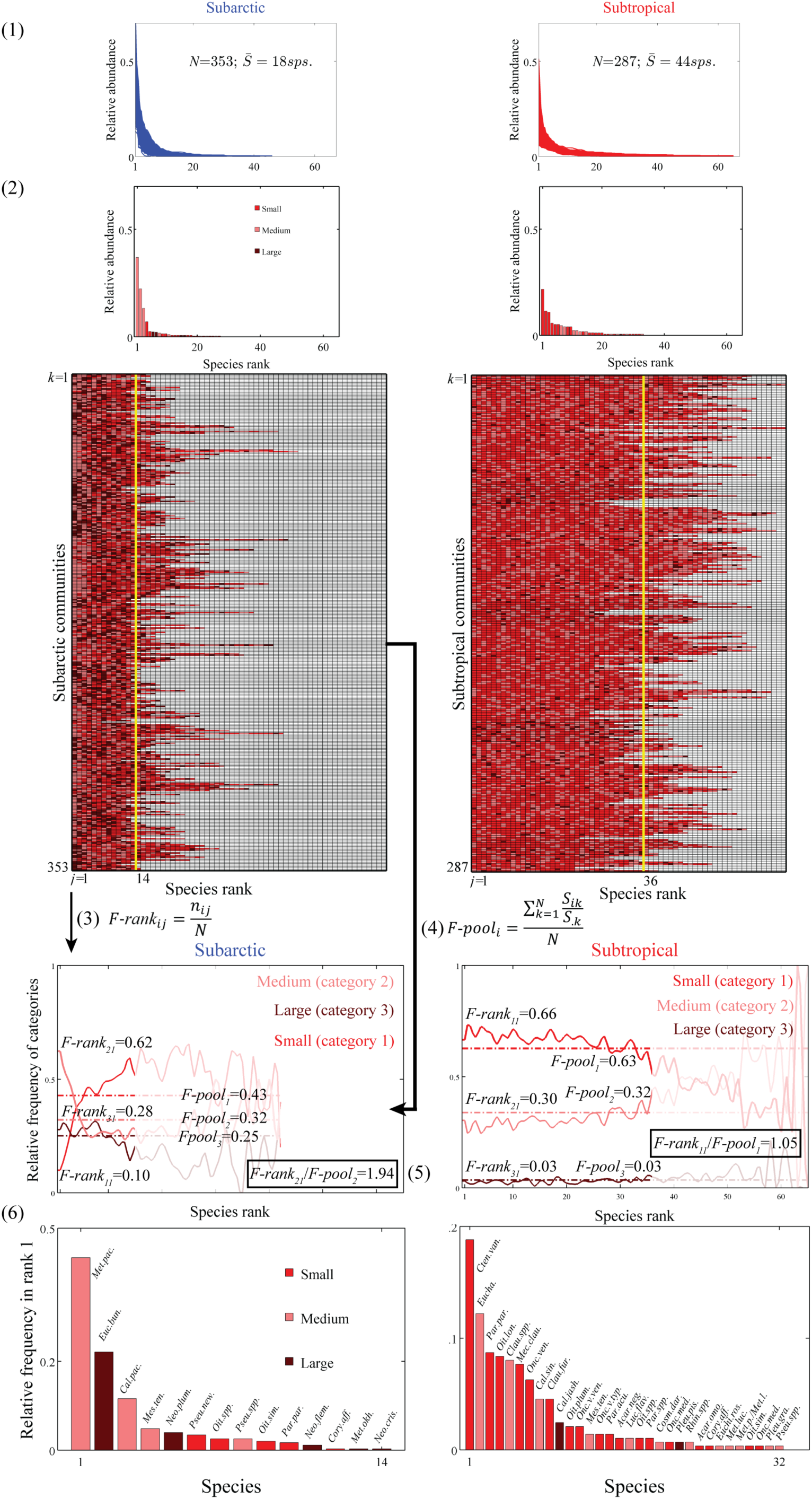
Schematic illustrating the method for the subarctic (left) and Subtropical (right) metacommunities: (1) for each local community, rank species from the most to the least abundant (i.e., constructing the RADs), thus producing *N* RAD curves for each metacommunity (*N*=353 local subarctic communities; *N*= 287 local subtropical communities); (2) for each local community, label the trait category for the species occupying each rank (e.g., is the species small, medium or large?); because local communities differ in number of ranks represented (i.e., differ in number of species), only the ranks represented in at least 80% of the local communities (e.g., yellow lines in checkerboard plots: ranks 1 to 14 in the subarctic metacommunity) are considered for visualization in the RADs; (3) for each metacommunity, quantify the relative frequency of categories in the species ranks (*F_rank_*, continuous lines in each panel) (e.g., in how many local subarctic communities the species occupying rank 1 are medium-sized?); (4) for each metacommunity, quantify the mean relative number of species belonging to each category in a community (*F_pool_*, horizontal dashed lines in each panel) (e.g., on average, how many species in a local subarctic community are medium-sized?); (5) test if the most-often-dominant category (e.g., medium size in subarctic communities) dominates more often than expected by chance (*F-rank_21_*/*F-pool_2_*>1) (e.g., medium-sized species dominate twice as often as expected by chance); *F-pool* further reflects niche overlapping related to legacy effects (e.g., no niche overlapping in subarctic communities versus overlapping towards small size in subtropical communities); (6) in order to explore the effect of species identity on the results, check the identity of the species occupying rank 1 in each metacommunity (i.e., few species occupy rank 1 and two species in each metacommunity dominate in more than 10% of the local communities): test the effect of species identity on *F-rank* through permutation of categories among the species occupying rank 1.

The relative frequencies of trait categories in the RADs (*F-rank*) show diverse dominance patterns among traits and regions, except for the feeding mode (Fig. 3 and Fig. 4). By contrast, regarding the mean relative frequencies of species with a trait category (*F-pool*) (i.e., as proxy for niche partitioning), in both regions, feeding mode shows strong over-redundancy, with most species being active (Table 1, *F-pool*; Fig. 4c, f dashed lines) while myelination and spawning strategy show a rather balanced partitioning between categories (Table 1, *F-pool*; Fig. 4g, j; h, k dashed lines). Only the mean relative frequencies (*F-pool*) for body size and thermal range differ regionally: subarctic communities show a more balanced partitioning among trait categories (Table 1, *F-pool*; Fig. 4a, i dashed lines) and subtropical communities show over-redundancy, with most species being small and warm-water (Table 1, *F-pool*; Fig. 4 d, l dashed lines). To depict the spatial pattern of trait category selection, we have also performed our analysis for 1°-1° grids (Fig. 5). The clearest pattern observed is that in northern communities, medium size, omnivory, active feeding, free-spawning, and cold are selected to dominate (Fig. 5a, b, c, e, f). Notice that sampling effort by grid is not homogenous (Fig. 3 and appendix S1) and samples in a grid may correspond to subarctic or subtropical communities depending on the position of water mass. Thus, we encourage gathering samples by water mass prior to applying the method and having similar sampling effort for the metacommunities as done in this study.

**Figure 3.**
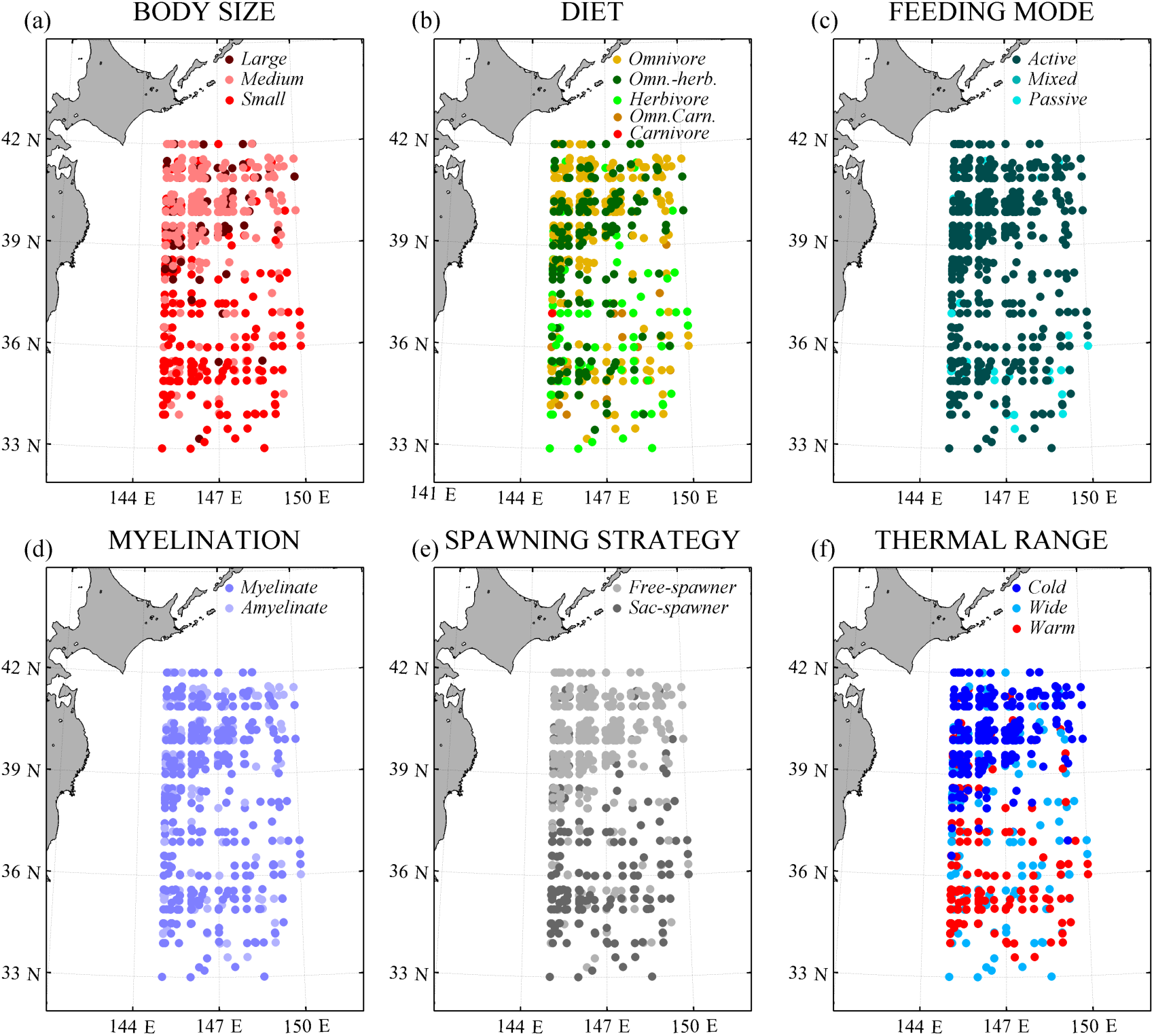
Geographic distribution of the samples, with colour corresponding, for each trait, to the trait category that occupies rank 1 in a community RAD. Notice that sampling is spatially (and temporally) heterogeneous (see appendix S1 for further information).

**Figure 4.**
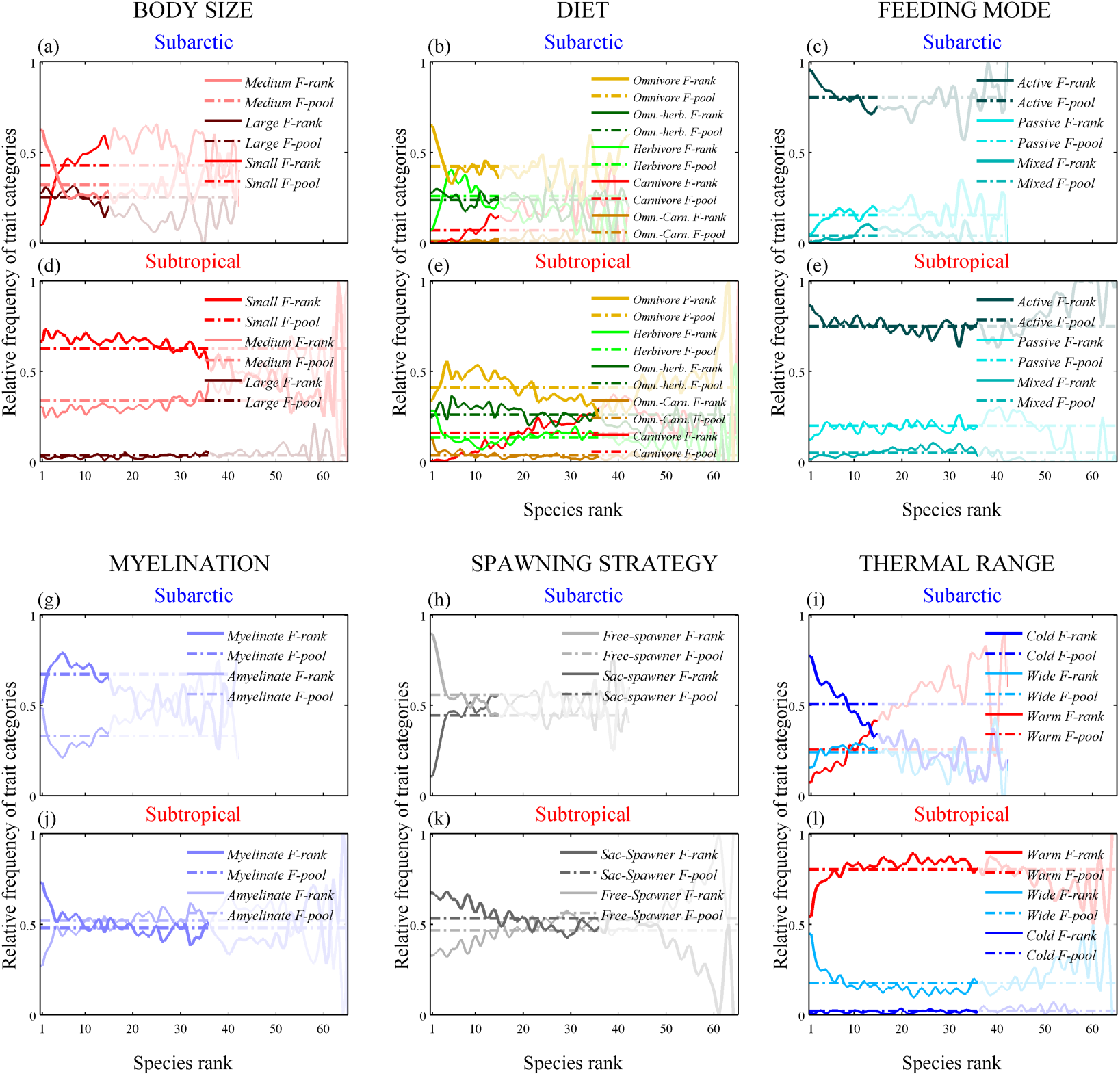
Relative frequencies of categories in the RADs of the two metacommunities (*F-rank*, continuous lines), and mean relative frequencies in the *N* species pools (*F-pool*, horizontal dashed lines). For each trait, the upper panel corresponds to the subarctic metacommunity (*N*=353), and the lower panel to the subtropical metacommunity (*N*=287). The shaded area in each panel corresponds to ranks that are underrepresented in the metacommunities and therefore are ignored. Categories in the legends are ordered from the greatest to the lowest *F-rank_1_*.

**Figure 5.**
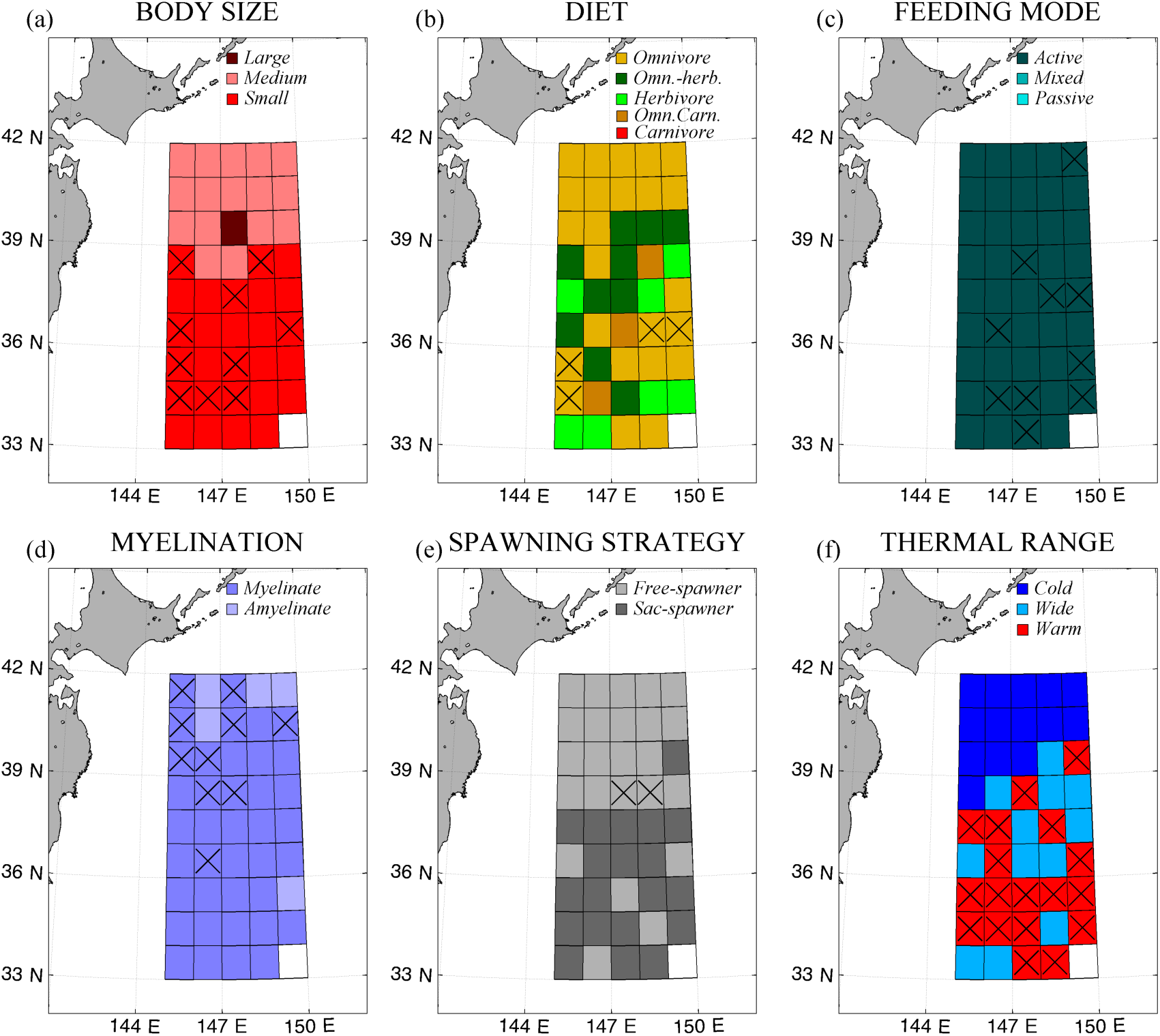
Results of the method applied to 1°-1° grid: colour of each grid corresponds to the trait category with the greatest *F-rank_1_*. Crosses mark grids where *F-rank_1_*/ *F-pool* ≤1, and thus we deem no selection of that trait category to dominate. Notice that sampling effort by grid is not homogenous and that samples in a grid may correspond to subarctic or subtropical communities depending on the position of the water mass. Thus, we encourage gathering samples by water mass prior to applying the method and having similar sampling effort for the metacommunities.

**Table 1.** Relative frequency of each category occupying species-abundance rank 1 (*F-rank_1_*) in the subarctic and subtropical metacommunities; mean relative frequency of species with each category (*F-pool*) and associated confidence intervals (*F-pool* _(C.I)_) for the most frequently dominant (greatest *F-rank_1_*) category (i.e., top category); and their ratio (top category *F-rank_1_*/*F-pool*_(C.I)_).

| TRAIT | Categories | *F-rank<sub>I</sub>* |  | *F-pool* (C.I.) |  | Top category<br>*F-rank<sub>I</sub>/F-pool* (C.I.) |  |
| --- | --- | --- | --- | --- | --- | --- | --- |
|  |  | Subarctic | Subtropical | Subarctic | Subtropical | Subarctic | Subtropical |
| Body Size | Small | 0.10 | 0.66 | 0.43 | 0.63<br>(0.62-0.63) |  | 1.06<br>(1.05-1.07) |
|  | Medium | 0.62 | 0.30 | 0.32<br>(0.31-0.33) | 0.34 | 1.94<br>(1.90-1.98) |  |
|  | Large | 0.28 | 0.03 | 0.25 | 0.03 |  |  |
| Diet | Herbivore | 0.07 | 0.28 | 0.26 | 0.13 |  |  |
|  | Omn.-Herb. | 0.28 | 0.25 | 0.24 | 0.26 |  |  |
|  | Omnivore | 0.65 | 0.34 | 0.42<br>(0.41-0.43) | 0.41<br>(0.40-0.42) | 1.53<br>(1.50-1.56) | 0.82<br>(0.81-0.84) |
|  | Omn.-Carn. | 0 | 0.13 | 0.01 | 0.03 |  |  |
|  | Carnivore | 0.003 | 0.003 | 0.07 | 0.16 |  |  |
| Feeding Mode | Active | 0.95 | 0.86 | 0.80<br>(0.79-0.81) | 0.75<br>(0.74-0.76) | 1.18<br>(1.17-1.20) | 1.15<br>(1.15-1.17) |
|  | Mixed | 0 | 0.01 | 0.04 | 0.05 |  |  |
|  | Passive | 0.05 | 0.12 | 0.16 | 0.20 |  |  |
| Myelination | Myelinate | 0.52 | 0.73 | 0.67 | 0.48 | 0.77<br>(0.76-0.79) | 1.52<br>(1.50-1.54) |
|  | Amyelinate | 0.48 | 0.27 | 0.33 | 0.52 |  |  |
| Spawning Strategy | Free-spawner | 0.89 | 0.33 | 0.56<br>(0.55-0.56) | 0.47 | 1.61<br>(1.58-1.63) |  |
|  | Sac-spawner | 0.11 | 0.67 | 0.44 | 0.53<br>(0.53-0.54) |  | 1.26<br>(1.24-1.28) |
| Thermal Range | Warm | 0.07 | 0.54 | 0.25 | 0.80<br>(0.79-0.81) |  | 0.68<br>(0.67-0.68) |
|  | Wide | 0.16 | 0.45 | 0.24 | 0.18 |  |  |
|  | Cold | 0.77 | 0.01 | 0.51<br>(0.49-0.52) | 0.02 | 1.52<br>(1.47-1.58) |  |

Body size and thermal range present the strongest regional differences in terms of both *F-rank* and *F-pool* (Table 1; Fig. 4 a, d; i, l). Medium, cold-water copepods dominate subarctic communities while small, warm-water copepods dominate subtropical communities (Table 1 greatest *F-rank_1_*; Fig. 4a, d). Medium instead of large species dominate in the subarctic communities, in accordance with Darmuth’s rule or the Energy Equivalence Rule ^27^; that is, for species using the same amount of energy, the smaller their size the more abundant they are. What is interesting is the higher proportion of small and warm-water species in the subtropical communities (Table 1, *F-pool*; Fig. 4 d, l), compared to more equal proportions (i.e. more balanced partitioning among body size and thermal range categories) in the subarctic communities (Table 1, *F-pool*; Fig. 4a, i). Thus, species niche packing is uneven with respect to body size and thermal range in subtropical communities, which agrees with the finding of over-redundancy towards the tropics recently reported for reef fishes ^24^. Furthermore, whereas in the subarctic, being medium-sized and cold-water confers greater relative fitness to a species and thus the advantage to dominate (Table 1 *F-rank_1_*/*F-pool*>1; Fig.4a, i), the effect of these functional traits (in the case of thermal range, the effect of a set of unknown functional traits that originate this biogeographic trait) on species fitness seems to be dampened towards the tropics; in the subtropics, the top-categories (i.e., small and warm-water) do not currently confer species an advantage to dominate (Table 1 *F-rank_1_*/*F-pool*∼1 and <1 respectively; Fig.4d, l; Fig. 5a, f). We further support our results with an additional analysis comparing relative total abundances of categories against the average abundance of species by category in both metacommunities (appendix S2). Our results point to an explanation for the previously reported greater species diversity despite non-concomitant greater functional diversity towards the tropics ^22,23^- a finding against expectations from classic niche theory.

The observed balanced partitioning between categories in the species pool and rank dominance of one category over the rest in the subarctic region are consistent with the expectations of classic niche theory of species coexistence that primarily explains that niche differences overcome differences in competitive ability (i.e., fitness differences) ^5^. Furthermore, in accordance with the storage effect that explains coexistence due to the temporal change of favourable conditions for the different species in a community ^8^, the competitive abilities of different categories vary seasonally in the subarctic region as environmental conditions change (appendix S3: seasonal signal of medium versus large size dominance). In contrast, in subtropical communities, the fact that most species are small and warm-water (Table 1 *F-pool*; Fig.4d, l) suggests that they coexist mainly because of small fitness differences, in agreement with modern coexistence theory ^8^. In fact, uneven niche overlapping towards the tropics has been reported for terrestrial plants ^22,28^ and reef fishes ^23,24^. This pattern agrees with the notions of stronger environmental filtering towards the tropics, evolutionary convergence, and niche conservatism.

We hypothesize that uneven niche overlapping facilitates high species diversity in subtropical communities through the following mechanism: species clumping in the best-adapted trait categories lowers their expected high relative fitness, facilitating invasibility for other less well adapted trait categories, and allowing those categories to dominate and persist in the communities. In the framework of the ‘time and area’ and ‘out of the tropics’ hypotheses, which assign the tropics as cradle and museum of species ^29–31^, we interpret our results as an evolutionary dependency of community structuring, with categorical traits conferring different fitness according to the environmental conditions at an initial time, followed by loss of that advantage as species overlap in the historically best-adapted categories (i.e., accumulated greatest fitness historically results in high F-pool of a category and eventually loss of greater fitness in current communities). This pattern and hypothesized mechanism have not, to our knowledge, been previously reported.

According to our hypothesis, we would expect that warm species have narrower pair-wise trait-based distances than cold species, and that species dominating more often tend to be more different (i.e., to have wider trait-based distances with the rest of species) in the subtropical set than in the subarctic set of communities. To test this hypothesis, we have computed the pair-wise trait-based distance among species in the space of a multiple correspondence analysis (MCA) applied on the five categorical functional traits following a previous study ^32^ (see MCA biplots in appendix S4; Fig. 6a) and compared these by thermal category (i.e., cold versus warm species). No significant difference exists between warm and cold species (Wilkoxon rank-sum test p=0,92) and dominant species do not show greater average pair-wise distances than the rest in the subtropical communities (Fig. 6a). These results do not support our hypothesis. Yet, our main results show that traits affect selection differently and thus distances with same weight for all traits may not be pertinent to test our hypothesis. Alternatively, we focused on body size as meta-trait for which contrasting patterns between the two sets of communities have motivated our hypothesis. We have compared the body size distribution of cold and warm species respect to the body size distribution of dominant species in subarctic versus subtropical communities (Fig. 6b). The results indicate that warm species are significantly smaller than cold species (Wilkoxon rank-sum test p=0,001) and have a narrower size distribution (Fig. 6b). What is more important is that body size of the most dominant species in warm waters (i.e., wide species denoted by light-blue stars with size corresponding to dominancy frequency) correspond to size out of the 1^st^ and 3^rd^ quartiles of the warm species size distribution (i.e., red data points out of the red boxplot depicting warm species size distribution) and thus far different from the average size of warm species (Fig. 6b). In contrast, the most dominant species in subarctic waters are cold water ones (dark blue stars), and the most dominant one has a body size that fits the median body size of cold species. Thus, body size distributions support our hypothesis that dominant species in subtropical waters tend to be more different from the rest of species which clump in a narrower niche range, as defined by body size, than in subarctic waters. This hypothesis rests to be further tested with other datasets.

Our results also support the hypothesis of increasing neutrality (i.e., similar fitness among species) towards the tropics ^33^. Yet, we cannot rule out the possibility that finer trait differences and/or intraspecific variability play more important roles in structuring and supporting diversity in subtropical communities than in subarctic communities. A meta-analysis of terrestrial animals suggests narrower niche breadths at lower latitudes mediated by richness (i.e., greater specialization with greater richness) ^34^ which would potentially support finer trait differences towards the tropics. Also, greater intraspecific genetic diversity in the tropics has been recently reported for terrestrial mammals and amphibians ^35^. Future observational studies quantifying intraspecific variability could therefore advance understanding of community structuring and diversity maintenance along gradients or across regions ^36,37^.

**Figure 6.**
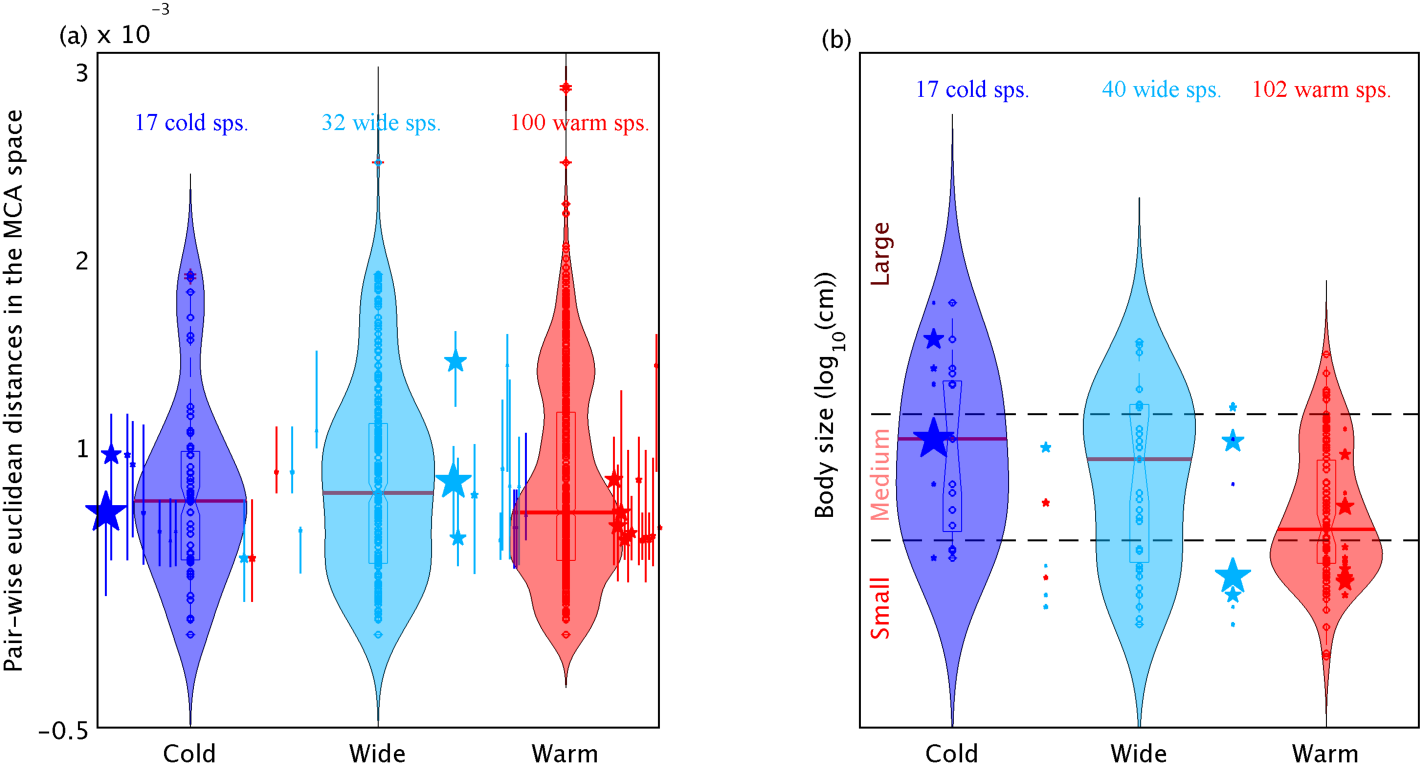
Violin plots (shadowed distributions) with superposed notched boxplots (vertical boxes), median (red horizontal lines) and single values (circles) illustrating (a) pair-wise trait-based species distances and (b) species nominal body size clustered by thermal range category. In (a), stars beside cold (blue) and warm (red) violin plots indicate the average pair-wise distance of dominant species (species occupying rank 1) in subarctic and subtropical waters respect to cold and warm water species respectively. Vertical lines associated with the stars delineate the 1^st^ and 3^rd^ quartiles of the pair-wise distances. In (b), stars beside the violin plots indicate the nominal size of dominant species; and horizontal dashed grey lines define the body size categories. Size of the stars denotes frequency of rank-1 occupancy, and colour matches the thermal category of the species.

Regarding the feeding-related traits, the dominant trait categories are the same in both regions: species with wider prey availability (i.e., omnivores), active search for their prey (i.e., active feeders), and more efficient predation escape reaction (i.e., myelinate) dominate (Table 1 greatest *F-rank_1_*; Fig. 3b, c, d; Fig. 4b, e; c, f; g, j). This empirical observation suggests three distinct hypotheses: (i) despite the different environmental conditions of such contrasting regions, trophic pressure (i.e., low prey availability and high predation pressure regardless of productivity level) and thus species interactions play a stronger role in determining the relative dominance of feeding traits; (ii) environmental filtering for those traits occurs at finer scales with similar conditions in both regions; or (iii) different environmental conditions in the two water masses select for the same trait categories due to selection through different co-varying factors (e.g., for feeding mode, known co-varying factors are prey availability, prey composition, predation risk and turbulence^38^). The current dataset does not allow us to investigate further, but we can compare with previous studies. Our results go against the expected dominance patterns for active versus passive feeding under different environmental conditions as mediated by relative food availability and predation pressure trade-offs ^39^, but agree with the global dominance of copepod active feeding ^26^ and myelination in the open ocean ^26,40^. Barton et al. ^50^ already anticipated the possibility of not observing biogeographical differences in feeding mode due to co-varying factors (i.e., predation risk-prey availability) but they expected changes at the seasonal scale. Seasonality of active versus passive feeding has been recently reported with community weighted mean indices ^26,41^. We, however, find no evidence for it in our dataset (appendix S3), although this finding should be taken with caution given our imbalanced sampling effort across seasons (appendix S1). Regarding myelination, our results of dominance in both water masses corroborate previous observations^40^. Lenz ^40^ suggests an advantage for myelinate copepods in environments with high predation pressure due to their enhanced escape response, but also in environments with low food availability due to their energy saving regarding impulse-related ion channels.

However, differences remain in the contributions of feeding-related traits between the two regions: a greater average contribution of carnivore species to the subtropical species pool (Table 1 *F-pool*; Fig.4b, e), and of myelinate species to the subarctic species pool (Table 1 *F-pool*; Fig.4g, j), respectively. The former agrees with the stronger latitudinal pattern of species richness for carnivores than for lower trophic levels reported by Hillebrand ^42^, whose meta-analysis included wide size ranges in each trophic level. Moreover, our results are in line with the distribution of some carnivore copepods in the global ocean^43^ and with the findings of greater relative abundance of carnivore copepods in the subtropics across the Atlantic Meridional Transect^44^. The greater prevalence of carnivore copepods in subtropical communities may be related to lower productivity in those waters that may confer ecological advantage to copepods specialising on food other than phytoplankton^43^ or it could also be related to greater prevalence of small copepods and thus more potential preys; yet, these rest as hypotheses and the underlying mechanisms remain unknown. The greater contribution of myelinate species in subarctic communities can be attributed to stronger predation risk from visual predators ^40^ in those waters, one of the most important fishing grounds in the North Pacific ^45^. It is also interesting that while feeding mode shows strong saturation towards active species in both regions (Table 1 *F-pool*; Fig.4c, f), the frequency is rather balanced for myelination (Table 1 *F-pool*; Fig.4g, j). Assuming that *F-pool* represents integrated effects of past selection and speciation, we interpret that, unlike feeding mode, myelination may not strongly affect species selection.

We observe a balanced frequency of the two spawning strategies in both metacommunities (Table 1 *F-pool*; Fig.4h, k). Thus, we assume weak selection regarding this trait, as previously suggested by Kiørboe and Sabatini ^46^. Yet, free-spawning species contribute slightly more to the species pool (*F-pool*=0.56) and tend to dominate in the subarctic communities (*F-rank_1_*/*F-pool*=1.61). In contrast, sac-spawning species contribute slightly more to the species pool (*F-pool*=0.53) and tend to dominate in the subtropical communities (*F-rank_1_*/*F-pool*=1.26) (Fig. 3e; Fig.4h versus k; Fig. 5e). Sac-spawning may be more affordable in warm waters due to thermal reduction of developmental times ^47^ and thus of maternal risk. It may also be more advantageous due to greater predation risk for free eggs in subtropical communities, given their more frequent copepod carnivory (*F-pool*= 0.16 versus 0.07) (Fig.4e versus b).

Finally, focusing on the identity of the species occupying rank 1 rather than on their trait category, we find that few species dominate in each metacommunity (i.e., 14 and 32 respectively) (Fig. 7). Furthermore, among those species, the relative frequency of rank 1 occupancy is highly unbalanced: only 2 of 14 species in the subarctic (Fig. 6 a-c, g-i) and 5 of 32 species in the subtropical communities (Fig. 6 d-f, j-l) occupy rank 1 in half of the communities (see list of species occupying rank 1 and frequency occupancy in appendix S5). A sensitivity analysis based on permutation of categories among the species reveals that in subarctic communities *F-rank_1_* is sensitive to the species identity (appendix S6). Several non-exclusive potential causes include: (1) other important traits that were not considered in this study, (2) differences in dispersion and immigration, and (3) neutral processes such as stochastic demographic drift that affect more strongly rare species, thus tending over time to support common species.

**Figure 7.**
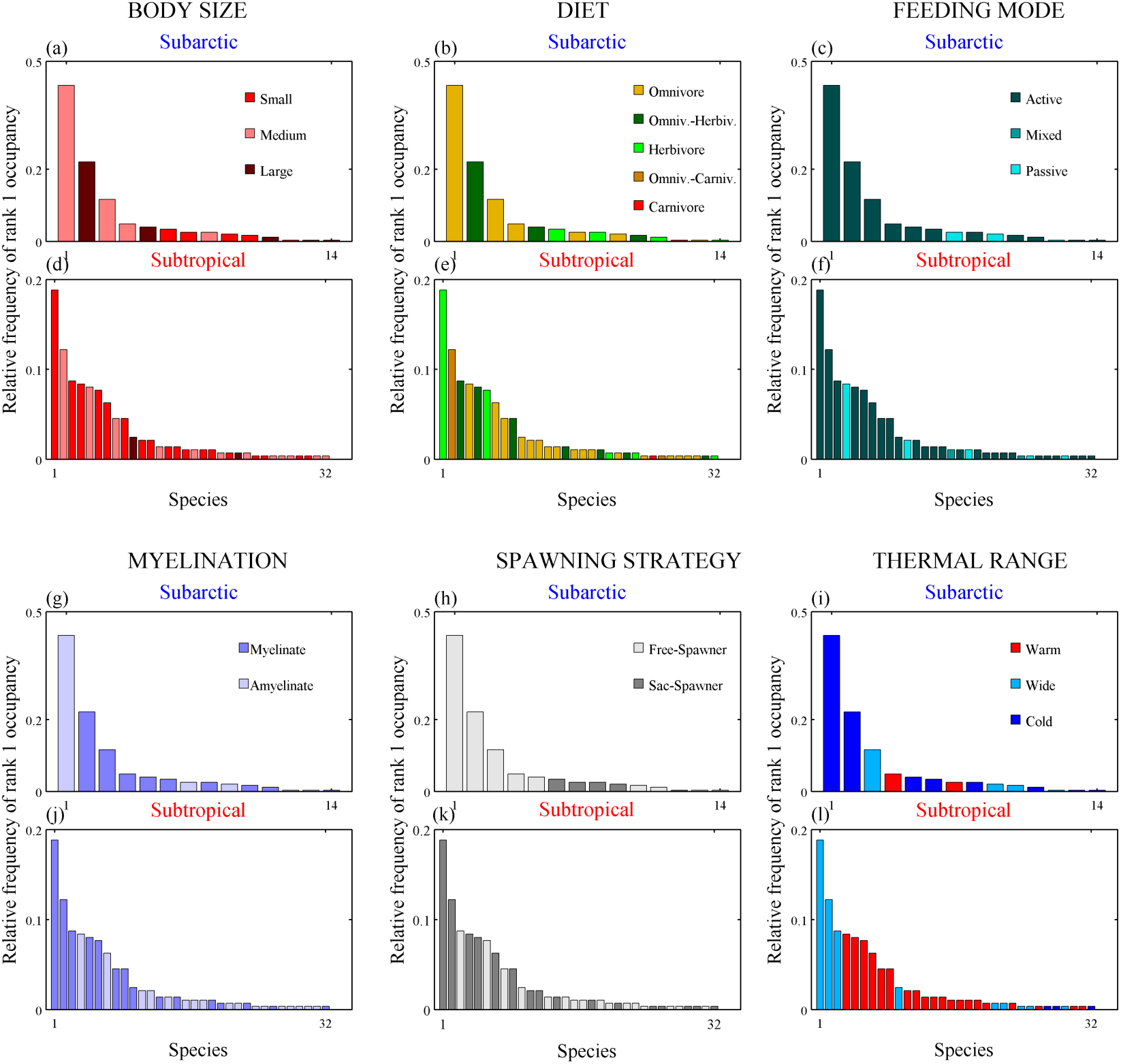
Relative frequency of rank 1 occupancy for each dominant species (14 species in the subarctic metacommunity and 32 species in the subtropical metacommunity), with colours indicating their categories.

Our new method, combining the RADs approach with trait information for the species occupying each rank in each local community, reveals that trait effects on copepod community structuring vary among traits and across regions (Fig. 4; Fig. 5).

Importantly, the shape of RAD has been theoretically linked to niche partitioning ^6,48^. As such, our approach allows distinguishing current selection of trait categories (*F-rank_1_* ∼ dominancy) from legacy effects such as integrated effects of immigration, past selection, adaptation and speciation (*F-pool* ∼ accumulation of species belonging to a certain category in the species pools). These findings would have been masked based on *integrated* functional diversity indices (see comparison with CWM in appendix S7). Nevertheless, our results must be taken with some caveats. The data, as all mesozooplankton net data, are biased towards larger copepods due to mesh retention limitations for the smallest abundant copepods. As stressed by Brun *et al.* ^26^, we neglected the contribution of abundant small passive feeding taxa such as Oithona spp. Nevertheless, *F-pool* of active feeding indicates that on average, ¾ of species identified in the communities were active, and presence detection is less prone to error of such small abundant copepods than are their abundance estimates due to net-mesh strainer effect. Also, the categorisation of species by traits is prone to error mainly due to lack of in situ observations of plankton behaviour and interactions. Here, we focused on copepods and neglected other competitors and predators of copepods such as gelatinous zooplankton, other macrozooplankton and fish. Finally, the nature of our dataset (i.e., historical species counts translated into categorical traits without concomitant environmental information) does not allow us to investigate further, but we encourage others with more complete independent datasets to further test the hypotheses that we have proposed. Specifically, datasets describing environmental and/or spatial gradients will allow proposing mechanisms explaining the new patterns unveiled with our method.

## Conclusions

In summary, we have presented a novel method to explore how functional traits structure communities by comparing the frequencies of trait categories in the species-rank abundance distributions (RADs) of local communities versus the frequencies of species with those trait categories in the regional average species pool. This method approaches a distinction between current selection (i.e., the degree to which a category confers dominance in the metacommunity ∼ high *F-rank_1_*) versus legacy effects such as integrated effects of immigration, past selection, adaptation and speciation (i.e., accumulation of species belonging to a certain category in the species pools ∼ high *F-pool*). Applying this method, we have unveiled the diverse effects of functional traits on the relative fitness of species and thus community structuring, and variation of these effects across regions. Overall, we observe relative fitness differences among species with different trait categories in the subarctic communities, as expected by classic niche theory, but not in the subtropical communities. Our results suggest an evolution of the role of influential categorical functional traits (i.e., body size as meta trait and thermal range as representing the outcome of a set of unknown environmental responsive functional traits) on community structuring, with fitness differences initially conferred, as observed in subarctic communities, and later lost as species clump in the best-adapted trait categories, as observed in subtropical communities. This mechanism potentially explains greater species diversity towards the tropics despite greater functional redundancy. The simplicity of our method allows straightforward application to observational species-counts data across systems and regions with the potential to advance understanding of the fundamental rules of community structuring, diversity maintenance, and the latitudinal gradient of species richness.

## Methods

### Dataset

Copepod community data represent a subset of the historical ODATE collection ^49^, which consists of zooplankton samples collected from 1960 to 2000 without associated environmental information (see appendix S1 for details). In order to compare subarctic versus subtropical communities, we delimited an area (Latitude: 33°-42° North; Longitude: 145°-150° East) in which sampling effort was the most balanced for both metacommunities (appendix S1, Fig. S1.1). This area corresponds to the confluence of the Oyashio (subarctic) and Kuroshio (subtropical) western boundary currents of the North Pacific. The Oyashio current is very productive with cold, nutrient-rich water that sustains an important fishing ground ^45^, whereas production is low in the nutrient-poor surface waters of the Kuroshio current ^50^. Copepods were sorted into species except when recognition could only be determined to the genus or family level. We considered all developmental stages because the contribution of juveniles to the community differs substantially in both datasets (appendix S1, Fig. S1.2). We separated subarctic from subtropical communities by water-mass identification following the local thermal criterion ^51^, and corroborated this separation by comparison with two clusters based on species composition (appendix S1, Figure S1.3). The resulting dataset consisted of a set of 353 subarctic communities, and a set of 287 subtropical communities.

### Species-level categorical functional traits

The six categorical functional traits analysed in this study were: body size as meta-trait; diet, feeding mode and myelination as trophic traits; spawning strategy as secondary-production trait; and thermal range as biogeographic trait (Box 1). These functional traits were chosen for their ecological meaning^52^ and for their availability at species-level classification. We were able to classify the most abundant and common 159 taxa of our samples (appendix S5), including 136 species-level, 20 genus-level and 3 family-level taxa. Our dataset included the vast majority of the sampled organisms in the subarctic communities, whereas for subtropical communities it included around 85% of the sampled organisms (appendix S1; Fig. S1.4). In any case, the identity of dominant species did not differ from the original dataset except for a few cases of highly aggregated taxa in the original dataset (appendix S1; Fig. S1.5). Trait classification at genus or family level was based on the classification of only the species belonging to that taxon that were present in the study area. Trait classification of the 159 species can be found in appendix S5.

### Data analysis

In order to explore whether species-level categorical functional traits affect the abundance ranking of species in a metacommunity as expected by classic niche theory, we have developed a simple method of trait-labelled species-rank abundance distributions (RAD) (Fig. 1 and Fig. 2):

1. We rank the species within each local community from most to least abundant (i.e., dominant to rare), constructing the *N* community RADs in a metacommunity.
2. In each local community, we label the trait category for the species occupying each rank and display the distribution of trait categories in the RADs (e.g., is the species in rank 1 small, medium or large?)

Because local communities differ greatly in number of ranks represented (i.e., differ in number of species), we focus on the distribution of categories in the *j* ranks present in at least 80 % of the local communities constituting each metacommunity (i.e., ranks 1 to 14 in subarctic communities and ranks 1 to 36 in subtropical communities).

(3) For each metacommunity, we then quantify in how many local communities each category occupied each rank: relative frequencies in each rank (e.g., how often the species occupying rank 1 was small?) (*F-rank*):

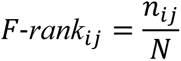

where for each functional trait, we count the number of local communities (*n*) in which the species belonging to the category (*i*) occupied each rank (*j*), separately for the subarctic metacommunity (*N*=353) and the subtropical metacommunity (*N*=287).

Conceptually, we interpret *F-rank* as representing current selection: greater frequencies towards the first ranks indicate the degree to which a category confers greater fitness than others in a metacommunity. Fitness, as the integrated ability to grow, survive and reproduce, is a primary determinant of the relative abundance of species.

(4) As reference for null selection, we calculate the mean relative frequency of species belonging to each category in a local community (e.g., on average, how many of the species in a community are small?) (*F-pool*):

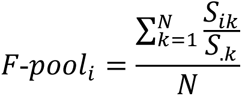

where *S_ik_* is the number of species belonging to category *i* in local community *k*, *S_.k_* is the total number of species in local community *k* (i.e., species richness), and *N* is the number of communities constituting the metacommunity. To determine the accuracy of *F-pool* as the mean value representing the average relative frequency of species belonging to a given category per local community, we estimate its 95% confidence intervals (C.I.) via 999 bootstraps of *F-pool_i_* (appendix S8).

Conceptually, we interpret *F-pool* as representing legacy effects such as integrated effects of immigration, past selection, adaptation and speciation: i.e., accumulation of species belonging to a certain category. In addition, *F-pool* is *per se* a proxy for species niche partitioning with respect to that trait (assuming that the trait affects species niche). An imbalance of *F-pool* among categories of a given trait reflects denser niche overlapping in the category with the greatest *F-pool*, and thus over-redundancy (e.g., most species in a community tend to be small). Alternatively, if *F-pool* is the same among categories (e.g., on average, in a community 1/3 of the species are small, 1/3 medium, and 1/3 large), niche partitioning with respect to that trait is evenly segregated (i.e., balanced). We further interpret balanced *F-pool* in both metacommunities to indicate weak past selection of species with respect to that trait. That is, no category has conferred integrated greater relative fitness to the species in any of the metacommunities.

(5) We test whether the top-category (i.e., category with greatest *F-rank_i1_*) occupies rank 1 more often than expected by chance, following a Bayesian probability reasoning: if *F-rank_i1_*/*F-pool_i_*>1, we assume that category *i* occupies rank 1 more often than expected by chance. Therefore, that category confers the greatest fitness to the dominant species in that metacommunity, or in other words, species having that category are currently being selected to dominate. Accordingly, *F-rank_i_* of that category shows a decreasing trend from rank 1 to lower ranks. Alternatively, if *F-rank_i1_*/*F-pool_i_*<1, we assume that the category is currently being selected against, and accordingly *F-rank_i_* shows an increasing trend from rank 1 to lower ranks. If *F-rank_i1_*/*F-pool_i_*∼1 we assume current neutral selection (deselection if *F-pool_i_* is high), and accordingly *F-rank_i_* shows a flat distribution along the ranks.

(6) Finally, we examine the sensitivity of results to species identity: we check the identity of the species occupying rank 1 in each metacommunity. We consequently conduct a sensitivity analysis based on permuting the categories among the species occupying rank 1 and re-computing *F-rank_1_* of the top-category to compare with its original estimate (appendix S6). The more variable the *F-rank_1_* estimates of 999 permutations are, the more sensitive are our results to the species identity (i.e., few species drive the observed results).

Additionally, in order to test contrasting mechanisms in subarctic versus subtropical communities suggested by our results, we calculated pair-wise trait-based distances among species and compared distributions between cold species (17), wide species (32) and warm species (100). Notice that these pair-wise distances were computed based on 149 species; for the other 10 species, their feeding strategy is unknown or is parasitic and thus was not considered in our analyses. We computed categorical distances on the space of orthogonal axes obtained from Multiple Correspondence Analysis (MCA) (see details on appendix S4). We further explored body size continuous distributions of the 159 species according to thermal range categories and contrasted body sizes of dominant species in subarctic versus subtropical communities in order to test our main hypothesis of trait-based community structuring.

## Code and data availability

Analyses were performed with MATLAB R2013b and a translation to R language of the new method is available at XXX together with the data generated and/or used in this study. The Odate collection is available at https://obis.org/dataset/e59f3fe1-8594-4dc4-81a2-58ddfa2a4026. For further information on data usage contact the metadata curator Dr. Tadokoro.

## Acknowledgements

We are grateful to the captains and crew of the research ships, to the researchers and technicians who collected and analyzed the samples. This work was supported financially by the Japan Science and Technology Agency, CREST grant number JPMJCR12A3 (P.I.: S.L.S). We thank Bingzhang Chen and Sergio Vallina for stimulating discussions.

### Box 1: Functional and biogeographic traits

#### Functional traits

##### Body size

Here, body size corresponds to the maximal total length of females as the species nominal size, retrieved from a local taxonomic key book ^53^. The three categories (i.e., small, medium and large) are defined according to the body-size tertiles of the log_10_-scale normal distribution of all species found in the ODATE collection (i.e., total length < 1.38 ≥ 3.71 mm) and corrected for the species shape (e.g., *Oithona setigera* would correspond to medium size class for its nominal size, yet it is classified as small for being elongated). In the case of broad taxonomic groups (i.e., genus), body size corresponds to the mean of nominal sizes of the species belonging to that group except for Clausocalanus spp. for which unclassified individuals tended to be larger and were classified as medium size. Body size is an easily measurable morphological trait that is considered a meta-trait because it has been reported to structure planktonic communities and determine their functioning in many ways via metabolic and predator-prey dependencies e.g., ^54,55–61^. We anticipate greater relative fitness, and therefore abundance dominance, of large taxa in subarctic communities and of small taxa in subtropical communities due to size-based competitive advantages related to water temperature and nutrient availability ^62–64^.

##### Diet

Diet corresponds to herbivore, omnivore, carnivore, and categories in between when explicitly recorded or when different bibliographic sources report one or the other (i.e., omnivore-herbivore and omnivore-carnivore). We consider that diet may structure communities via energy constraints (i.e., prey availability, individual prey value and energy expenses in capturing and handling the prey) ^65,66^.

##### Feeding mode

We classified copepod species as having active, passive or mixed feeding mode, following a recent compilation of copepod functional traits ^67^. For this trait, 12 taxa were excluded for being parasitic or because their feeding mode is unknown (appendix S5). The feeding mode entails trade-offs between energy expenses and gain, as well as between predation risk and chance of mating ^39^. Active feeding entails more risk but also more gain, while passive feeding has lower predation risk but also lower prey encounter ^68^. Because of these trade-offs, environmental conditions, especially turbulence, predation pressure, and prey quantity and quality (i.e., motility) are expected to affect the frequency of feeding strategies in the communities ^39^.

##### Myelination

We classified the taxa as myelinate and amyelinate based on the phylogeny of calanoid copepods ^40^. Copepods with myelin covering their nervous system are reported to have more precise escape reactions to predation ^69^. Lenz ^40^ proposes that myelinate copepods should dominate in waters with high predation risk (e.g., subarctic) because they have better capacity to escape. However, she also proposes that they may dominate in waters with low nutrient availability (e.g., subtropical) too, because myelinate axons require less energy than amyelinate axons, and because waters with low productivity are more transparent, increasing predation risk. Therefore, we anticipate greater fitness, and hence abundance dominance, of myelinate copepods in both metacommunities.

##### Spawning strategy

Spawning strategy distinguishes between copepods that release their eggs in the water column (free-spawn) or carry their eggs until hatching (sac-spawn). Free-spawning strategy consists of releasing many eggs in the water with low maternal investment per egg and low individual egg survival, whereas the sac-spawning strategy consists of producing fewer eggs per female unit biomass with the mother carrying them until hatching. Thus, sac-spawning implies greater individual egg survival ^70^ but greater predation risk for the mother ^46^. The two strategies have implications for energy allocation, with a trade-off between number of eggs and developmental time until hatching ^71^ and another trade-off between survival rates of individual females and their eggs ^46^. According to Kiørboe and Sabatini ^46^, because of those trade-offs, we expect both strategies to be equally fit and therefore none should dominate over the other.

#### Biogeographic trait

##### Thermal range

The thermal range corresponds to the regional biogeography of the taxa in relation to thermal conditions (e.g., some copepod species are primarily found in warm waters). According to bibliographic research, we classify the taxa as those found mainly in cold waters, warm, or with a wide distribution across the Northwest Pacific ^53^. We do not consider it as a trait *per se* but as an outcome of several functional traits that may confer a taxon the specialized ability to inhabit certain environments. Accordingly, we anticipate greater relative fitness and therefore abundance dominance of categories in their corresponding water masses (i.e., cold in subarctic and warm in subtropical communities).

## Appendix S1: Details of dataset construction and structure

We analysed a subset of the Odate collection (Odate 1994), which consists of zooplankton samples collected with vertical hauls of a NORPAC net (45-cm diameter and 330-µm mesh), from 150 m depth to the surface. Samples were collected by various Japanese research institutes from 1960 to 2000 and stored by the Japan Fisheries Research Agency. In order to compare subarctic and subtropical communities, we chose an area in which sampling was the most balanced for both sets of communities (Latitude: 33°-42° North; Longitude: 145°-150° East) (Fig. S1.1). This area corresponds to the confluence of the Oyashio (subarctic) and Kuroshio (subtropical) western boundary currents of the North Pacific. The Oyashio current is very productive with cold, nutrient-rich water that sustains an important fishing ground (Sakurai 2007), whereas production is low in the nutrient-poor surface waters of the Kuroshio current (Qiu 2001). Oyashio zooplankton biomass has a strong seasonality with accumulation around late spring, while Kuroshio zooplankton biomass remains lower having milder seasonality with two low peaks in spring and autumn (Yoo *et al.* 2008). There were no environmental data associated with this dataset; in addition, because samples were collected during the period 1960-2000, related satellite observations were scarce.

**Figure S1.1.**
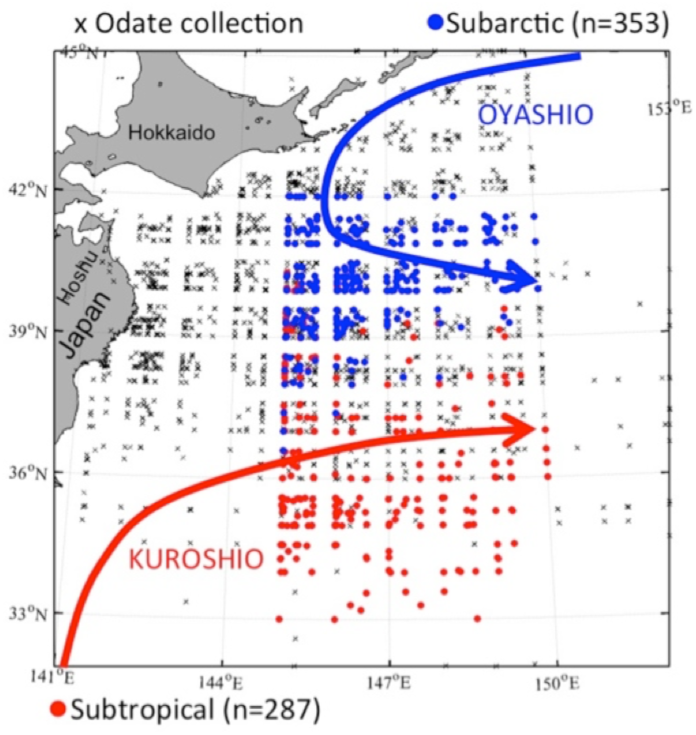
Sampling location, in the area 33°-42° North 145°-150° East where the Oyashio and Kuroshio western-boundary currents meet.

Copepods were sorted into species, except when they could only be identified to the genus or family level. We considered all developmental stages (i.e., copepodite to adult) because the contribution of juveniles to the community and their identification to the species level differed substantially by region. On average, 70% of organisms in the subarctic communities corresponded to juveniles versus 30% in the subtropical communities; yet, 80% of juveniles were classified to the species level in subarctic communities versus 30% in subtropical communities (Fig. S1.2).

**Figure S1.2.**
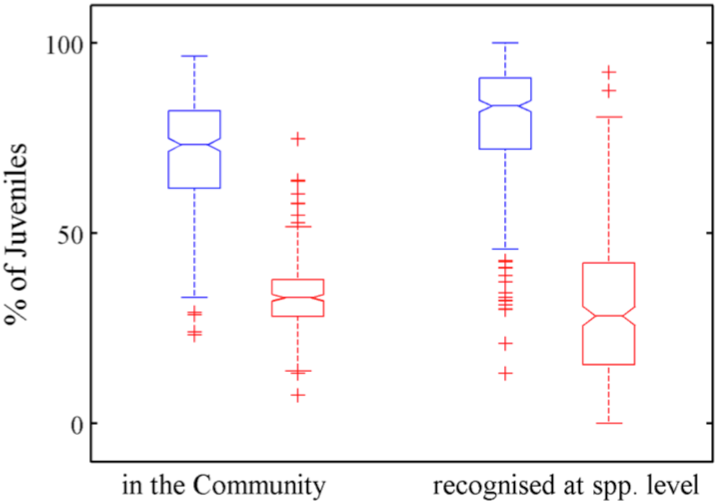
Boxplots depicting the distribution of percentage of juveniles in the communities and their recognition to the species level for subarctic (blue) and subtropical (red) communities.

Samples were first sorted as subarctic or subtropical based on hydrographic conditions of the Oyashio and Kuroshio currents in this area (i.e., 100-m depth temperature <5°C and 200-m depth temperature >15°C respectively) (Kawai 1972). Then, we corroborated the separation into two sets of communities by sorting the communities into two clusters according to the Hellinger-transformed species abundances and ordination by nonmetric multidimensional scaling (NMDS) (Legendre & Legendre 1998). The Hellinger transformation was applied to prevent disproportionate effects of rare species on the clustering. The two clusters were obtained by hierarchical clustering following Ward’s criterion. The two sets of communities matched for both classifications, except for two samples that were classified as subarctic according to the water mass but were determined as subtropical according to the species composition. We decided to keep the original hydrographic classification of the two sets of samples after inspection of sample ordination by NMDS on the Euclidean distances of the Hellinger-transformed abundances (Fig. S1.3a). We also inspected seasonality in the dataset (Fig. S1.3b-c). As usual in oceanography, winter was highly under-sampled in both sets of communities (Fig. S1.3c).

**Figure S1.3.**
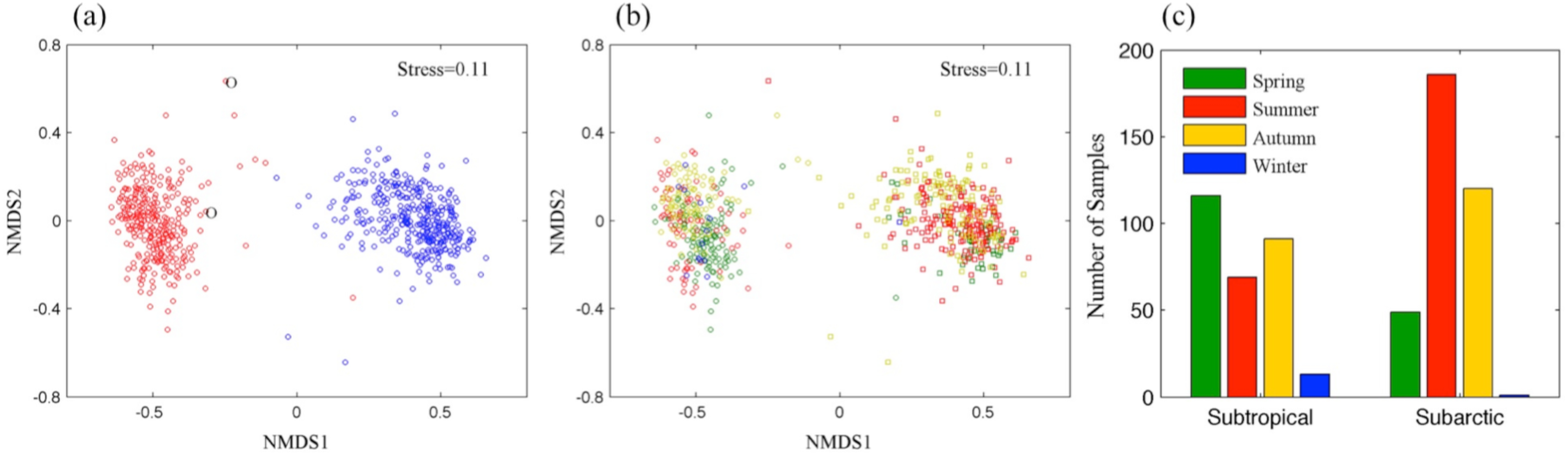
(a) Samples ordered by NMDS; blue circles correspond to the subarctic communities and red circles correspond to subtropical according to the clustering; superposed ‘O’ label corresponds to samples classified as subarctic according to water mass but subtropical according to the species composition. (b) Same as in (a), but colour corresponds to seasons (legend in subplot (c)); (c) Seasonal representation in the two sets of communities.

Based on bibliographic information, we managed to classify 159 taxa into six categorical functional traits (appendix S2). The 159 taxa corresponded to 136 species, 20 genera and three families (see the list in table appendix S2). Our dataset of 159 taxa included the vast majority of the sampled organisms in the arctic communities, and around 85% in the subtropical communities (Fig. S1.4).

**Figure S1.4.**
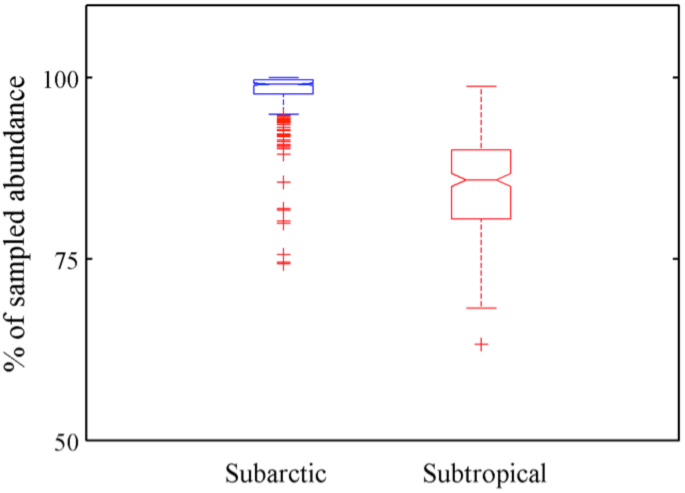
Boxplots depicting the percentage of sampled organisms that were retained in our 159-taxa analysis, in subarctic (blue) and subtropical (red) communities.

We tested whether this subsampling would affect the identity of the taxon dominating each community (the main focus of our study). The identity of the dominant taxon differed for one subarctic community and for 55 subtropical communities (0.2% and 19% of samples in each set) (Fig. S1.5), yet these corresponded to highly aggregated taxa (i.e., the order Calanoida and genera *Eucalanus* spp. and *Pleuromamma* spp.) in the original dataset.

**Figure S1.5.**
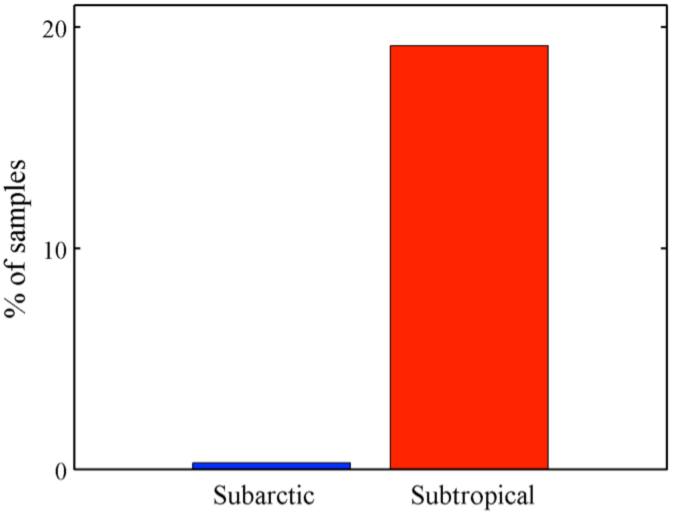
Percentage of subarctic (blue) and subtropical (red) samples for which the dominant species differs from the original dataset.

Samples were spatially heterogeneous, with the strongest sampling effort in the northwestern side of the area (Figure S1.6a). When diving the data into 1°-1° grids, the number of species classified in a community ranged on average from 15 to 50 species from north to south (Figure S1.6b).

**Figure S1.6.**
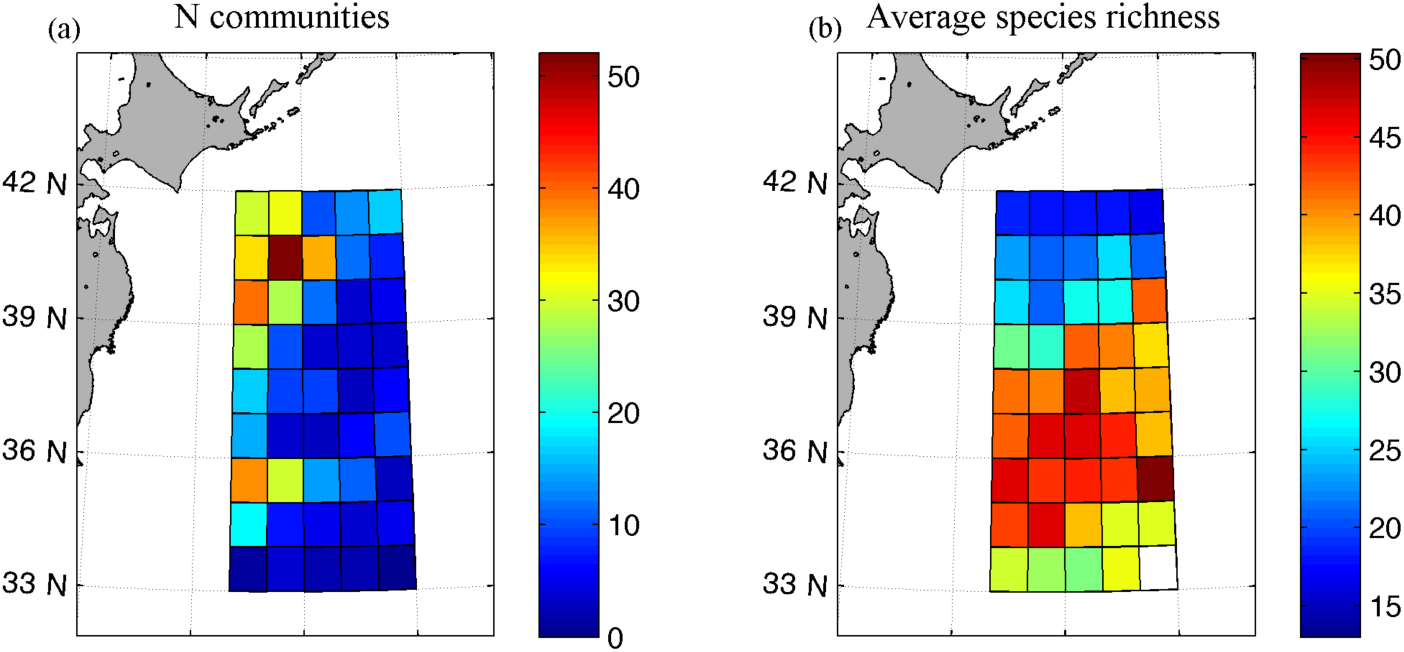
(a) Sample frequency and (b) average number of species in a community in each 1°-1° grid.

**Figure S1.7.**
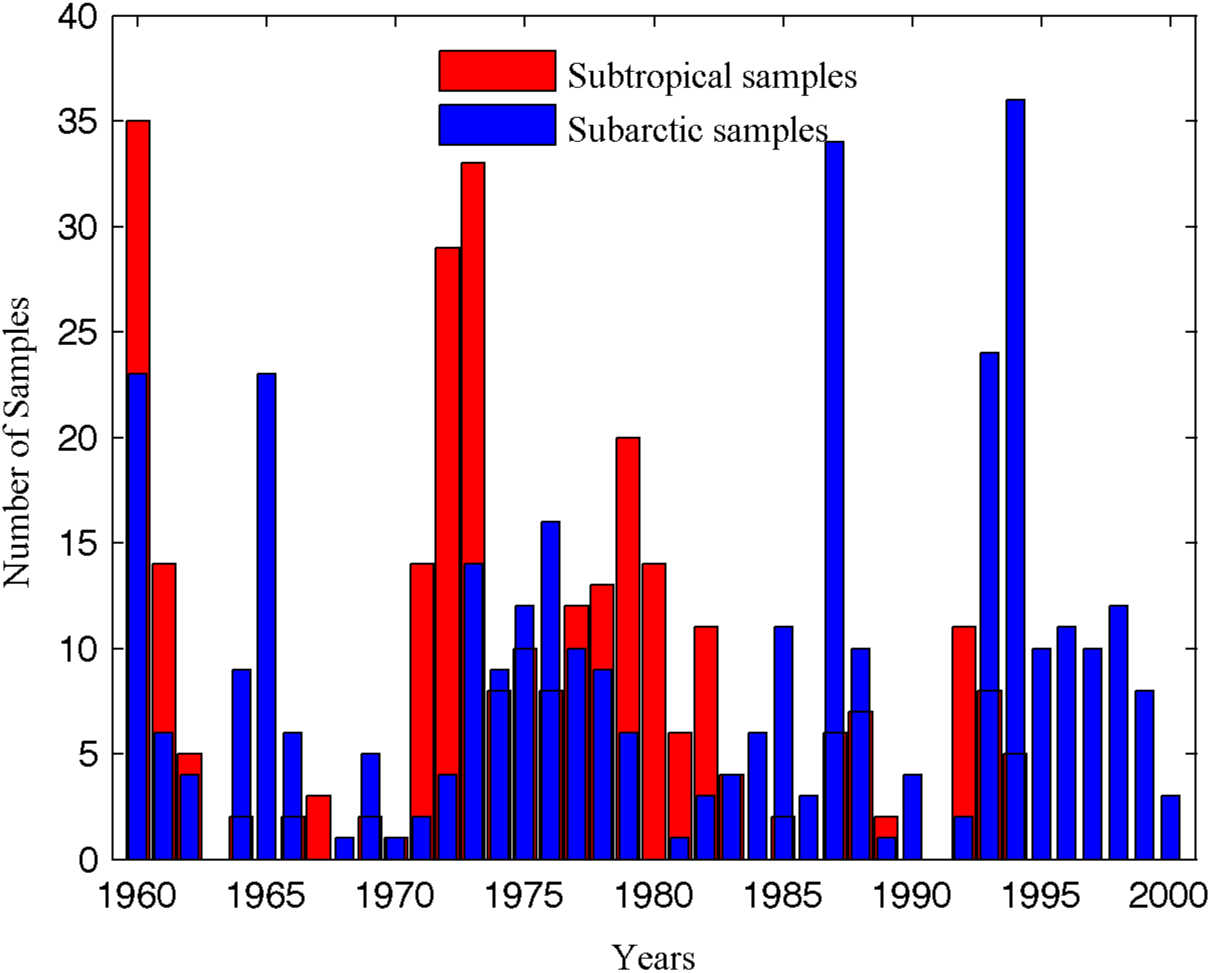
Sample frequency of the two water masses by year.

Samples were also temporally heterogeneous, with subtropical samples being more frequently sampled in the 70s and not sampled in the last five years of the series. This interannual heterogeneity is due to financial availability and increasingly targeting sampling lines corresponding to more productive areas, as the main purpose of the Odate collection was to monitor prey available for fish in the Japanese fisheries grounds (Sugisaki 2006).

## Appendix S2: Comparison of normalized average abundances of species by category with the relative abundance of categories in the communities

Because our method is based on species ranks, neglecting abundance information, we support our main results with an additional analysis. We compare the relative abundance of a category in a community versus the relative mean abundance of a species with that category in the community respect to the null expectation of even relative abundance of species in the community (i.e., equal relative fitness). That is, we plot for each category *i*:

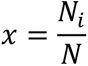

where *N_i_* is the abundance of copepods with a category *i* in a community, and *N* is the total abundance in that community; versus

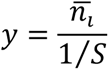

where 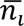 is the relative mean abundance of the species with category *i* in the community and *S* is the number of species in the community so that 1/*S* corresponds to the expected abundance of the species following the null expectation of even abundances of the species in the community (Figure S2.1). Furthermore, for each category we calculated the percentage of communities in which *y* was greater than 1 and the 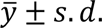. A high 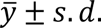 and the prevalence of *y*>1 suggests that the category confers greater relative fitness to the species (table S2.1). This analysis corroborates our main results. In the subarctic, being medium-sized and cold-water confers greater relative fitness to a species and thus the advantage to dominate (in most communities, medium-sized and cold-water *y* is greater than 1; Figure S2.1, table S2.1); the effect of functional traits on species fitness is dampened towards the tropics: in the subtropics, the top-categories (i.e., small and warm-water) are the ones most represented in the community (greater *x* and 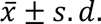; figure S2.1, table S2.1) and do not currently confer species an advantage to dominate respect to the other categories (*y*∼1 in figure S2.1 and low % in table S2.1).

**Figure S2.1.**
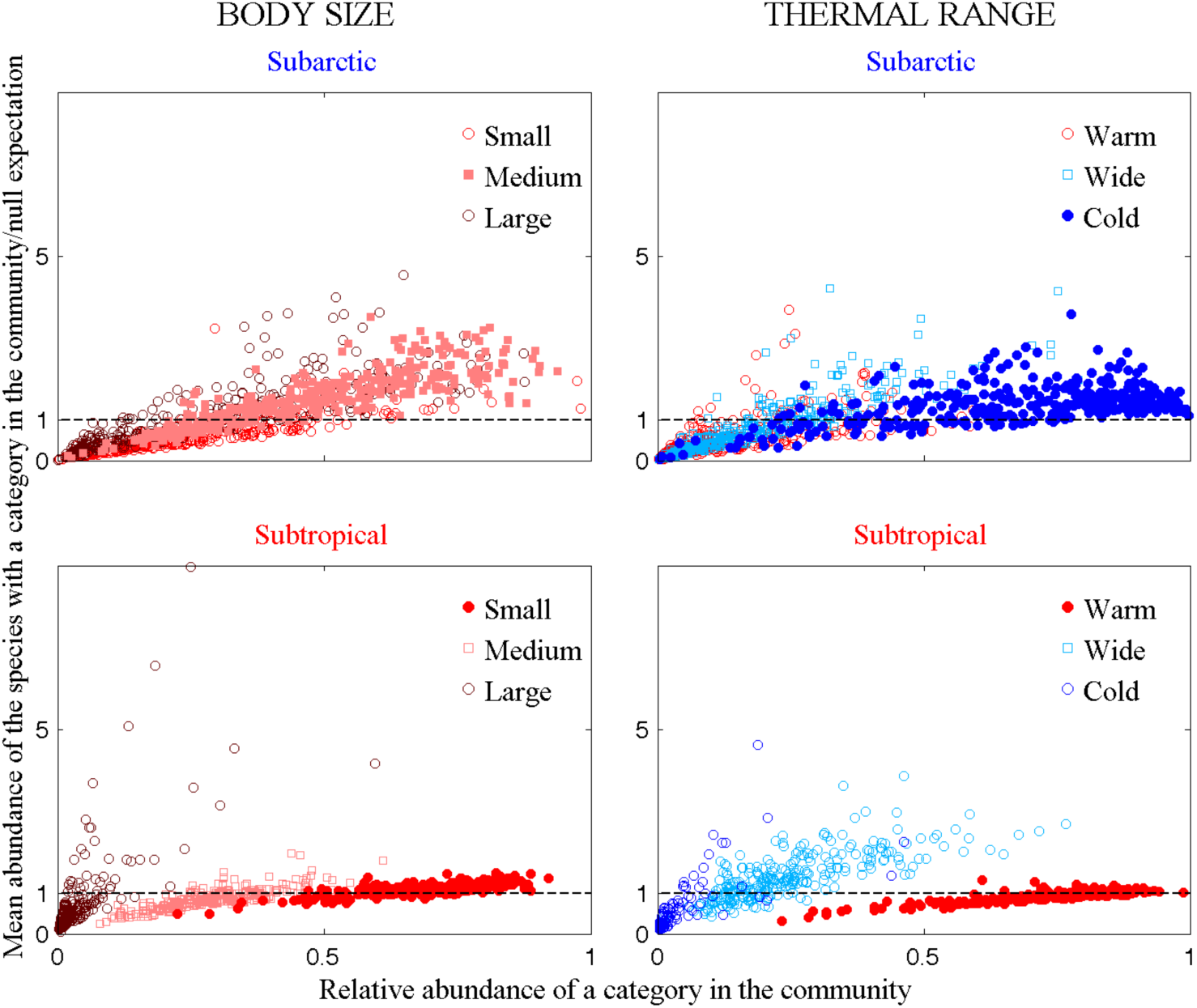
Relative abundance of a category in the community (*x*) versus the mean abundance of species with a category in the community/null expectation (*y*).

**Table S2.1.**
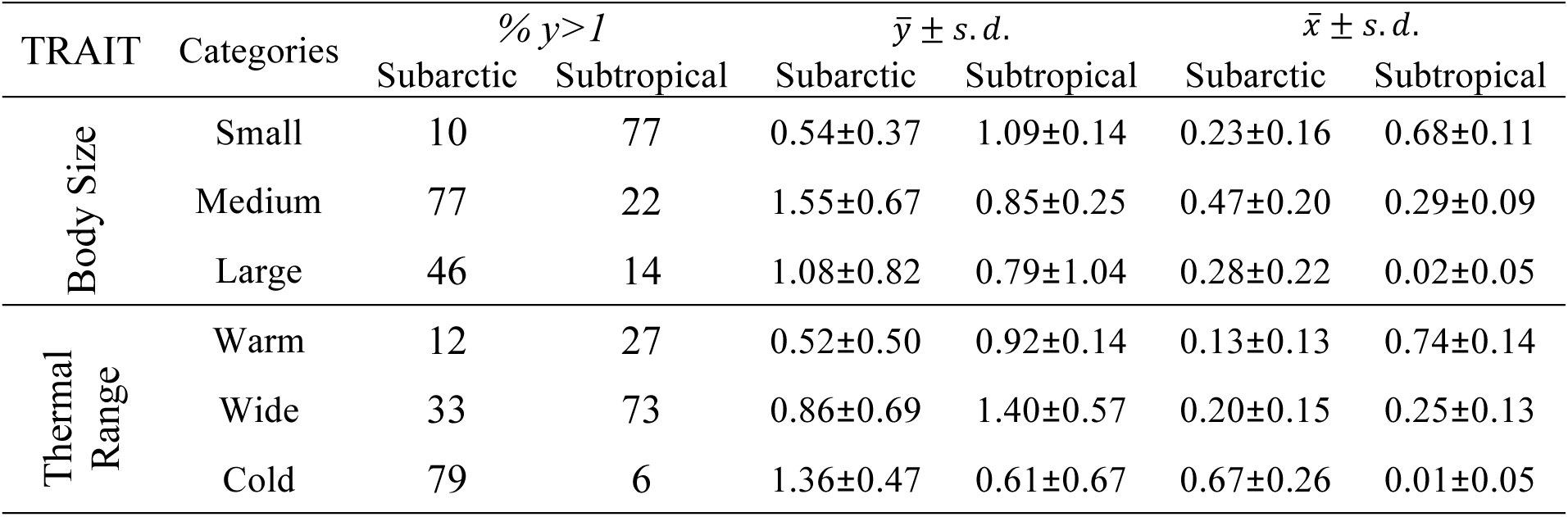
Percentage of communities in which the mean abundance of the species with a category in the community respect to the null expectation is greater than 1 (%*y*>1) and mean value and standard deviation of *y* (*ȳ* ± *s. d*.) and *x* (*x̄* ± *s. d*.) in the two sets of communities

## Appendix S3: Seasonality of *F-rank* and *F-pool*

We explored the seasonal pattern of *F-rank* and *F-pool*. We should keep in mind that summer is overrepresented in the subarctic set, and winter is represented in neither of the sets of communities. Regarding *F-rank_1_*, we observe seasonality for body size and diet in both subarctic and subtropical communities. In subarctic communities, there is greater rank 1 occupancy of large species in spring (brown continuous line at rank 1 in Fig. S3.1 a) in contrast with almost consistent rank 1 occupancy of medium species in autumn (pink continuous line at rank 1 in Fig. S3.3 a), and greater rank 1 occupancy of omnivory-herbivory and herbivory in spring than in the other seasons (dark green and green continuous lines at rank 1 in Fig. S3.1 d). In subtropical communities, in spring, contrary to the other seasons, we observe selection of small (red continuous line at rank 1 in Fig. S3.1 e) and herbivore species to dominate (green continuous line at rank 1 in Fig. S3.1 d). Regarding *F-pool*, it is rather stable among seasons except for thermal range in subarctic communities. We observe greatest frequency of cold versus warm species in spring (horizontal lines in Fig. S3.1 i) and lowest in autumn that could be explained by stronger inmigration of subtropical species to subarctic waters in autumn (dashed lines in Fig. S3.3 i).

**Figure S3.1.**
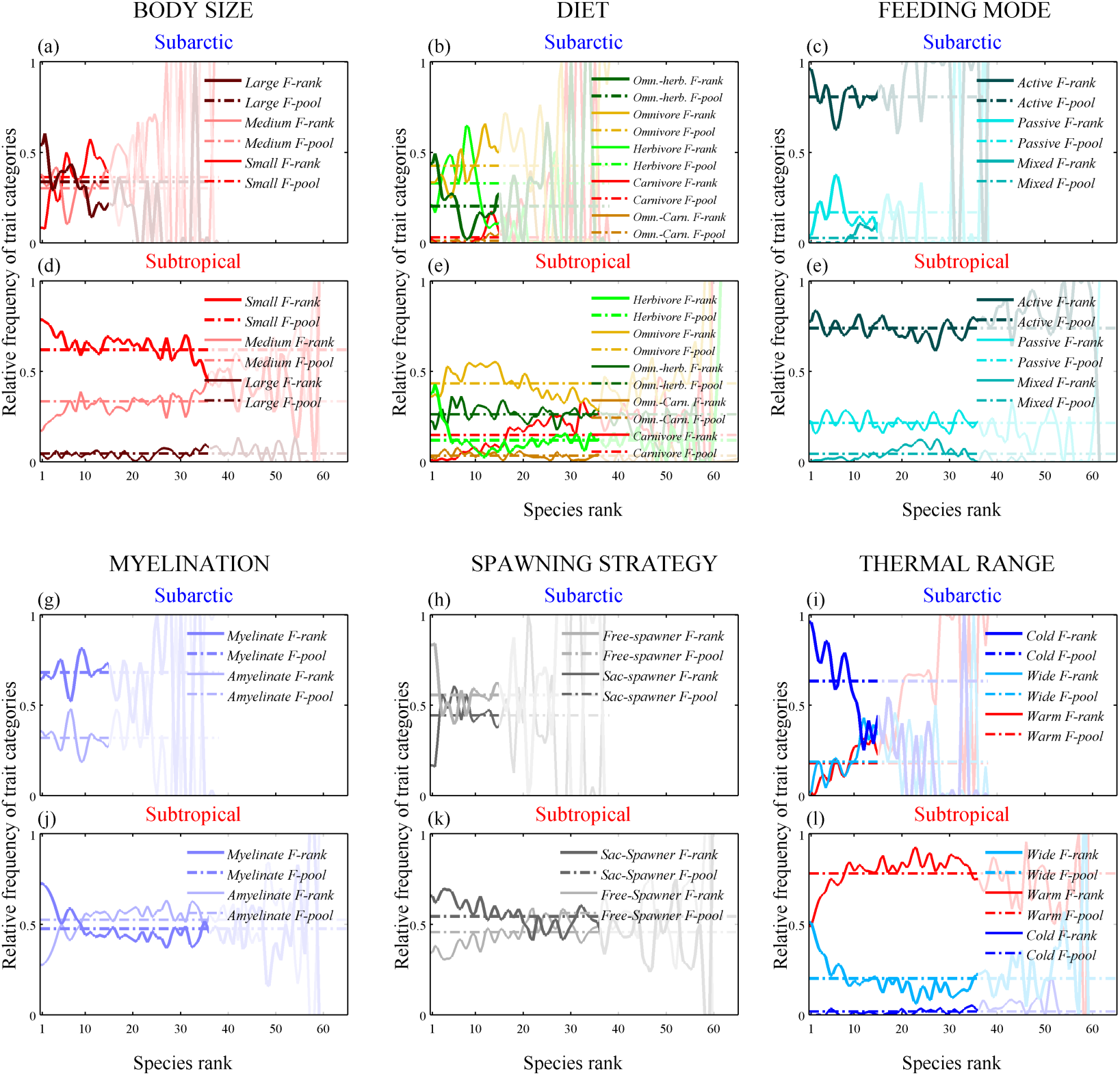
Same as figure 2 in the main text but for communities only sampled in spring: 48 subarctic communities versus 116 subtropical communities.

**Figure S3.2.**
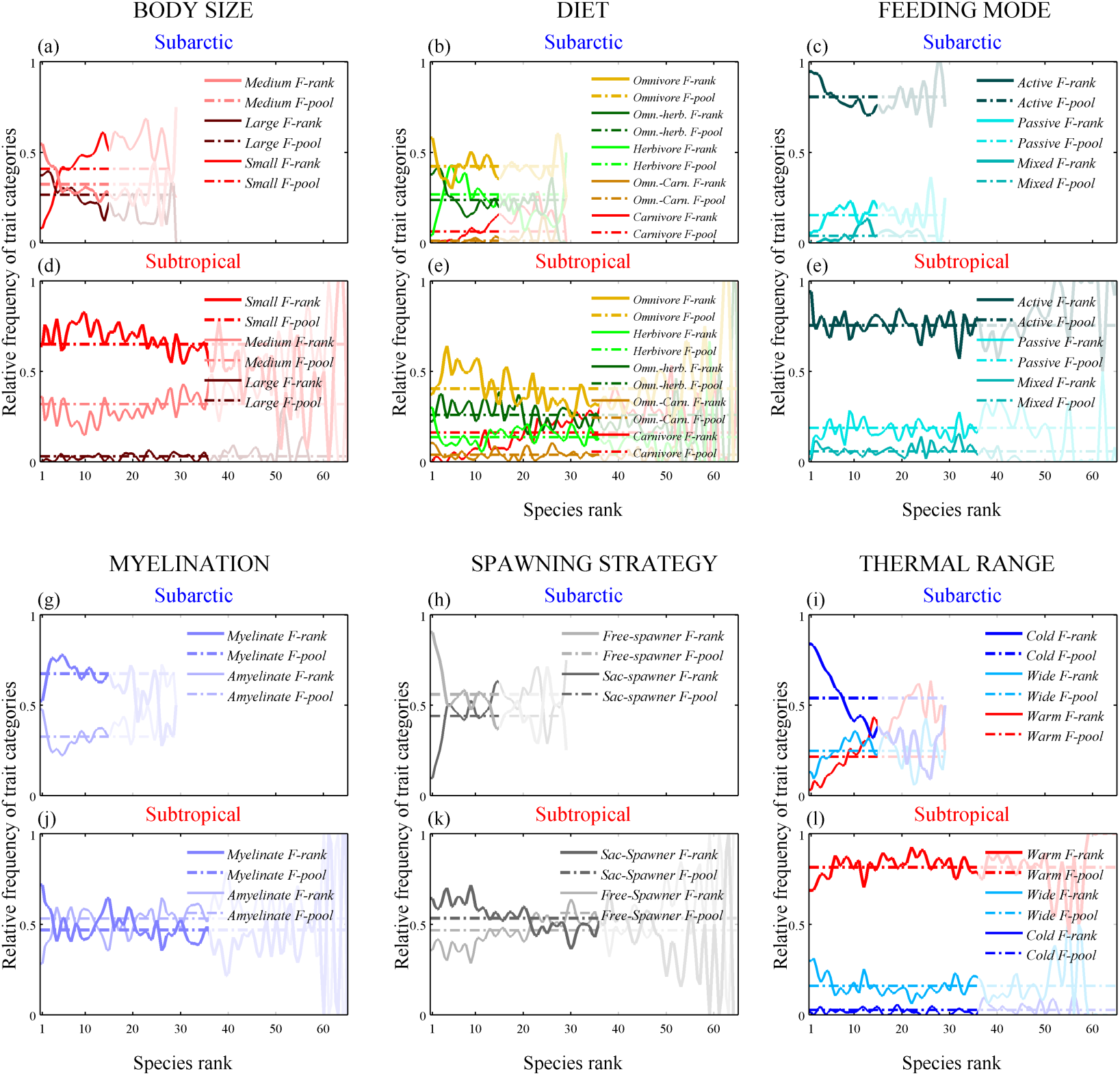
Same as figure 2 in the main text but for communities only sampled in summer: 185 subarctic communities versus 67 subtropical communities.

**Figure S3.3.**
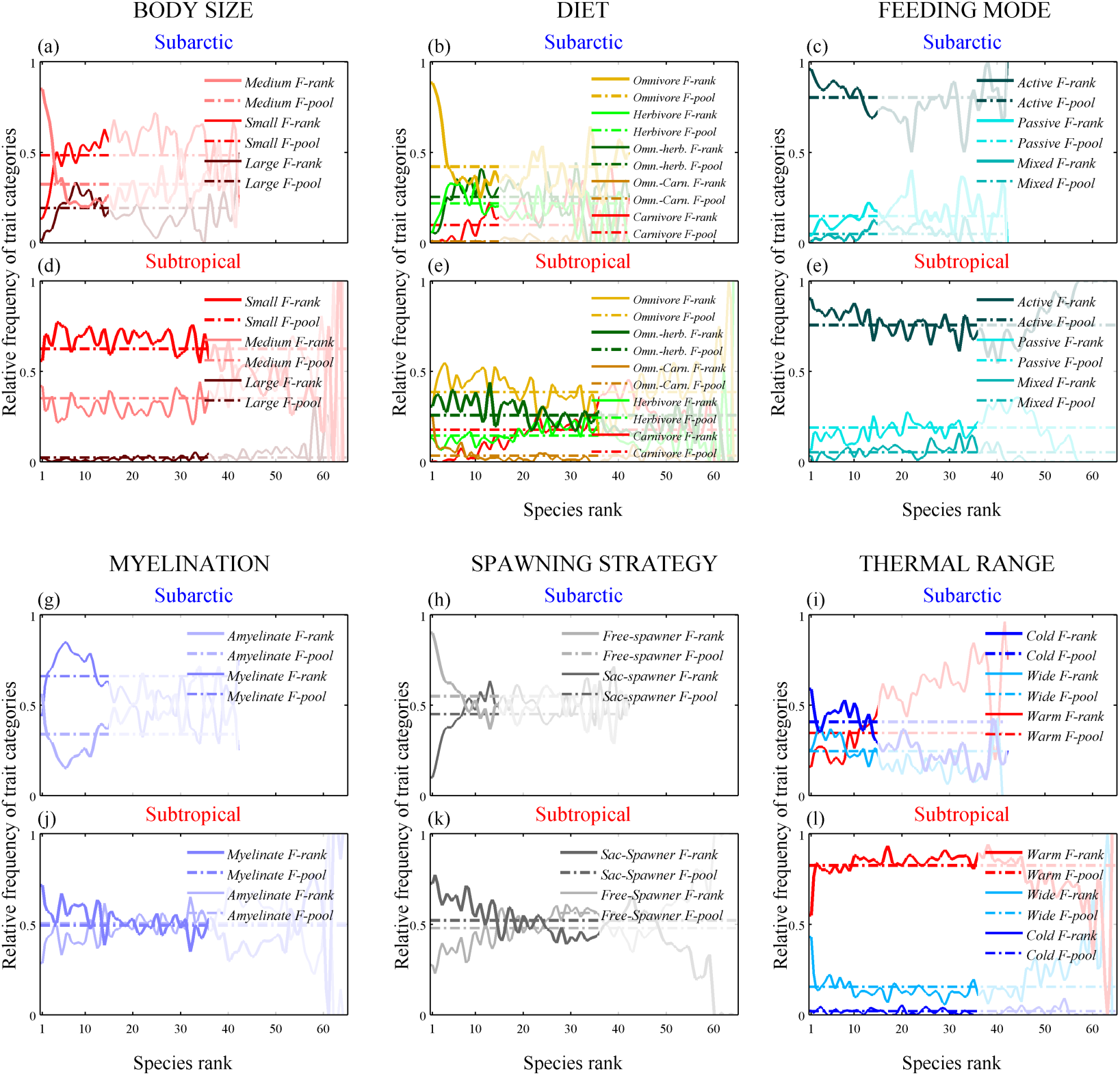
Same as figure 2 in the main text but for communities only sampled in autumn: 119 subarctic communities versus 91 subtropical communities.

## Appendix S4: Multiple Correspondence Analysis (MCA) of the species traits

In order to calculate the trait-based distances, among warm species, cold species and wide species respectively (Figure 5a), we applied Multiple Correspondence Analysis (MCA) on the five categorical functional traits (Figure S4.1). We computed the species pair-wise Euclidean distances in the MCA orthogonal space (Fig. S4.1) according to the Kaiser-Guttman criterion ^1^: 5 eigen vectors representing 73% of the total variance.

**Figure S4.1.**
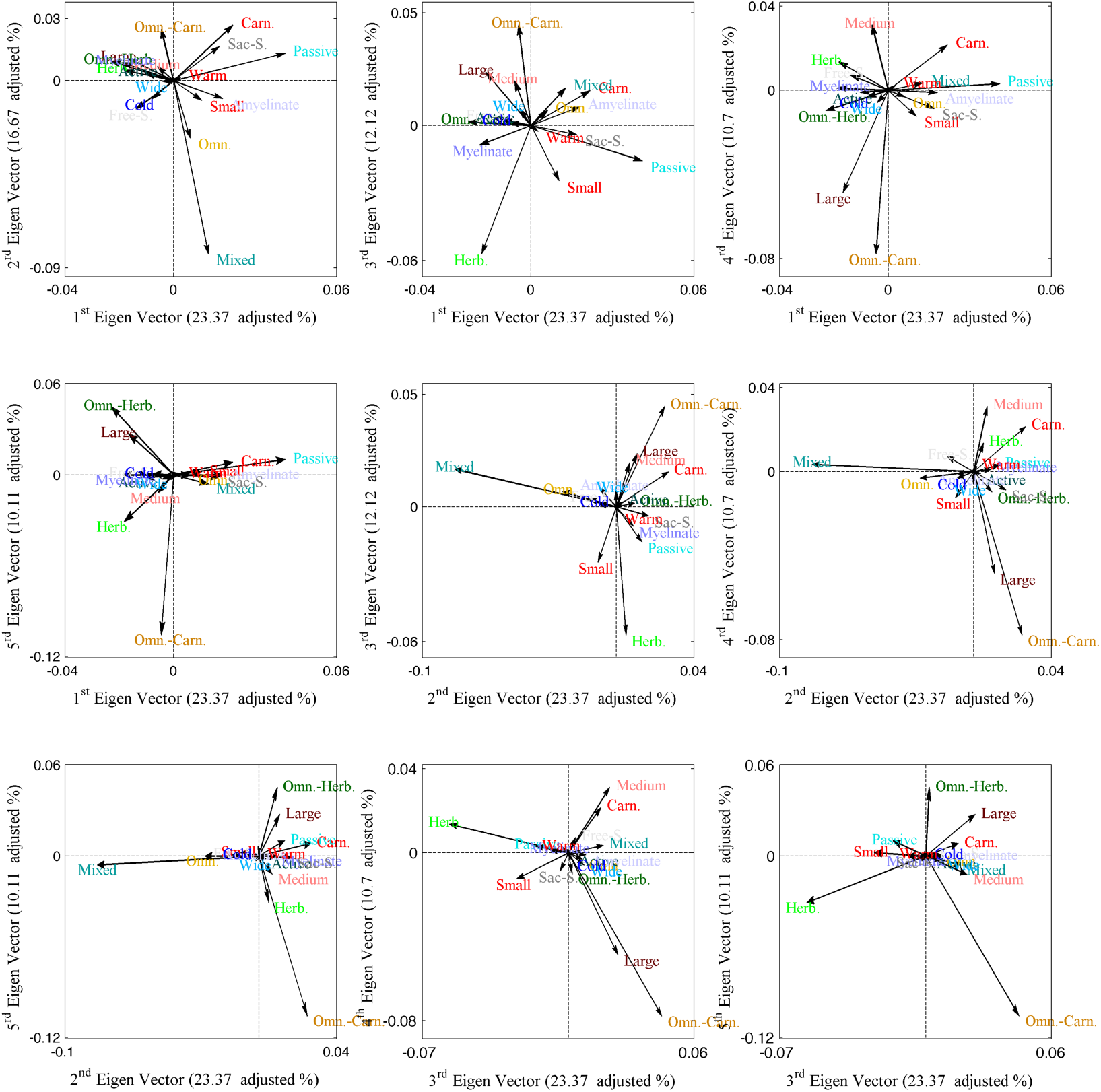
Structuring of the trait categories in the orthogonal spaces of the 1^st^ 5 eigen vectors of the MCA, with warm, cold and wide categories predicted as additional trait (i.e., not participating in the ordination but positioned afterwards in the orthogonal spaces).

## Appendix S6: Sensitivity of the analysis to species identity

Species identity is a potential concern because relatively few species occupy rank 1 in both sets of communities (14 in the subarctic and 32 in the subtropical set), and therefore trait frequencies are to some extent driven by species identity. Furthermore, among those species, the relative frequency of rank 1 occupancy is highly unbalanced: 2/14 species in the subarctic and 5/32 species in the subtropical communities occupy rank 1 in half of the communities. In order to explore the sensitivity of our analyses to species identity, for each trait we randomly permuted the trait categories (*n*=999) of the 14 and 32 species occupying rank 1 for the subarctic and subtropical metacommunity respectively, and for each iteration we recomputed *F-rank_1_* of the top category. We compared the randomly generated values (histograms) with the actual value of the top category *F-rank_1_* (black line) (Fig. S6.1). The greater the variability of the randomly generated values (i.e., greater range and flatness of the distribution), the greater the sensitivity of our analysis to species identity. We found that subarctic communities present a great sensitivity (Fig. S6.1). Also, the position of the generated values with respect to the actual top category *F-rank_1_* reflects the tendency of that trait category to characterise species that occupy frequently rank 1 versus those that rarely do it in the set of communities: the more the histogram is positioned to the left of the actual *F-rank_1_*, the more the species that frequently dominate rank 1 are characterised by the top category, and conversely, if the histogram is to the right of the actual *F-rank_1_* the top category tends to characterise the species that dominate less frequently.

Except for myelination in subarctic communities and for omnivory and warm-water in subtropical communities, the rest of top categories tend to characterise the most often dominant species (Fig. S6.1). Finally, for the subtropical communities, if the mode of the generated distribution matches *F-pool*, the frequency of categories in the species occupying rank 1matches the mean frequency of categories in the community (i.e., body size, feeding mode, myelination and thermal range).

**Figure S6.1.**
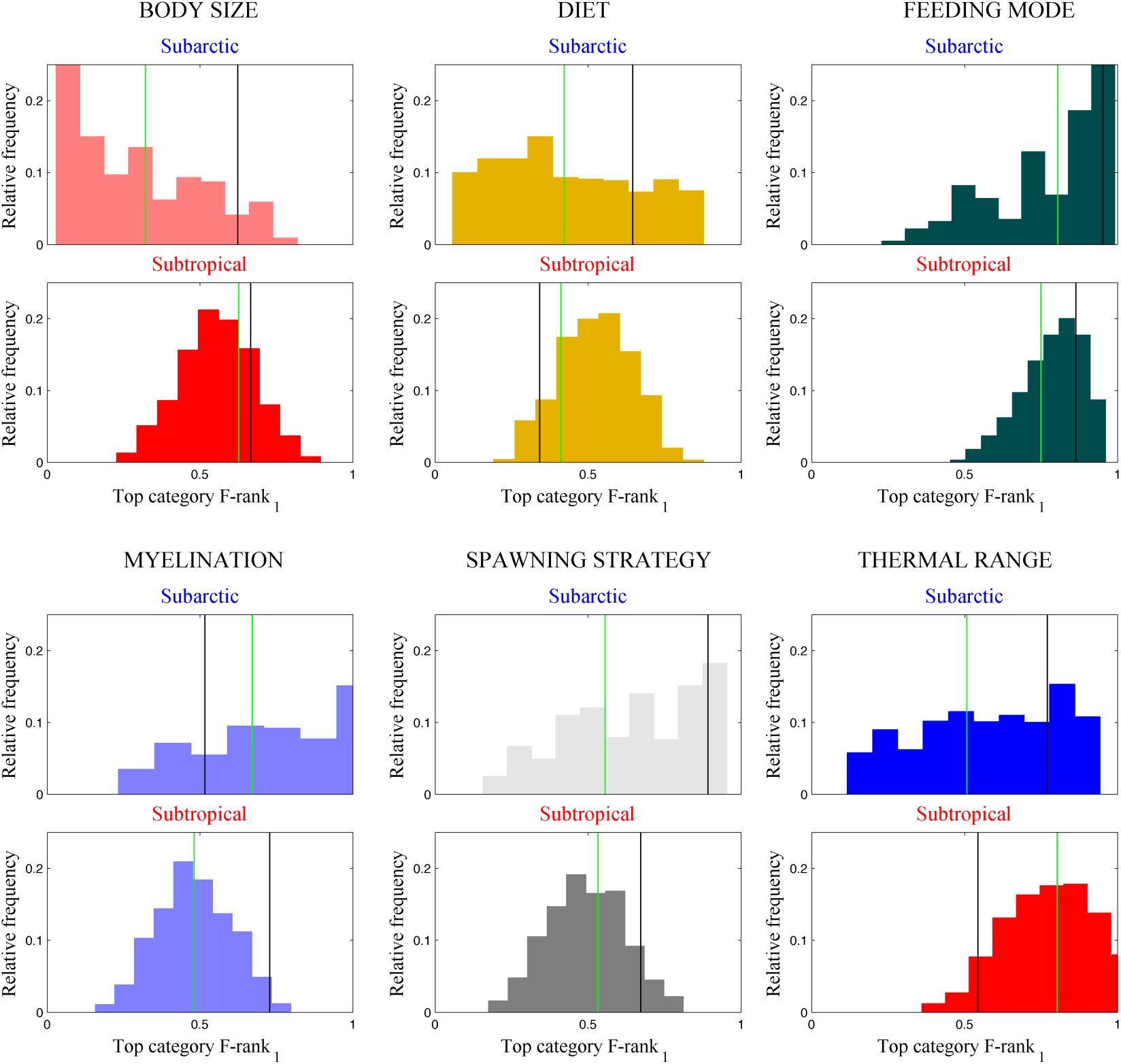
Top category *F-rank_1_* (black line) with respect to its estimates in 999 random permutations of the dominant-species trait categories (histograms), and to *F-pool* (green line).

## Appendix S7: Comparison of our method versus the community weighted mean (CWM) approach

We argue that community-integrated measures do not inform whether the observed changes in trait composition are due to changes in the proportion of species with certain trait values (i.e., relative richness) or due to changes in the abundance of species (i.e., relative abundance). This issue is critical if we aim to understand the role of traits on community structuring (i.e., species relative fitness) and to test whether a trait affects species abundance dominancy.

In order to demonstrate that community-integrated measures are not suitable to test if a trait category affects species relative fitness and therefore dominancy, we compared our method with its analogous of community-integrated measures: the community weighted mean (CWM). Specifically, we compared rank 1 occupancy of each body size category with the body size category determined by the CWM body size. The CWM body is calculated as:

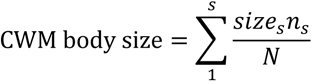

where size*_s_* is the nominal body size (i.e., maximal total length of females from a taxonomic key book) of species *s*, *n_s_* is the abundance of species *s* in the community and *N* is the total abundance in the community. For similar values of CWM body size, we observed very different species rank abundance distributions (RADs) (Fig. S7.1).

**Figure S7.1.**
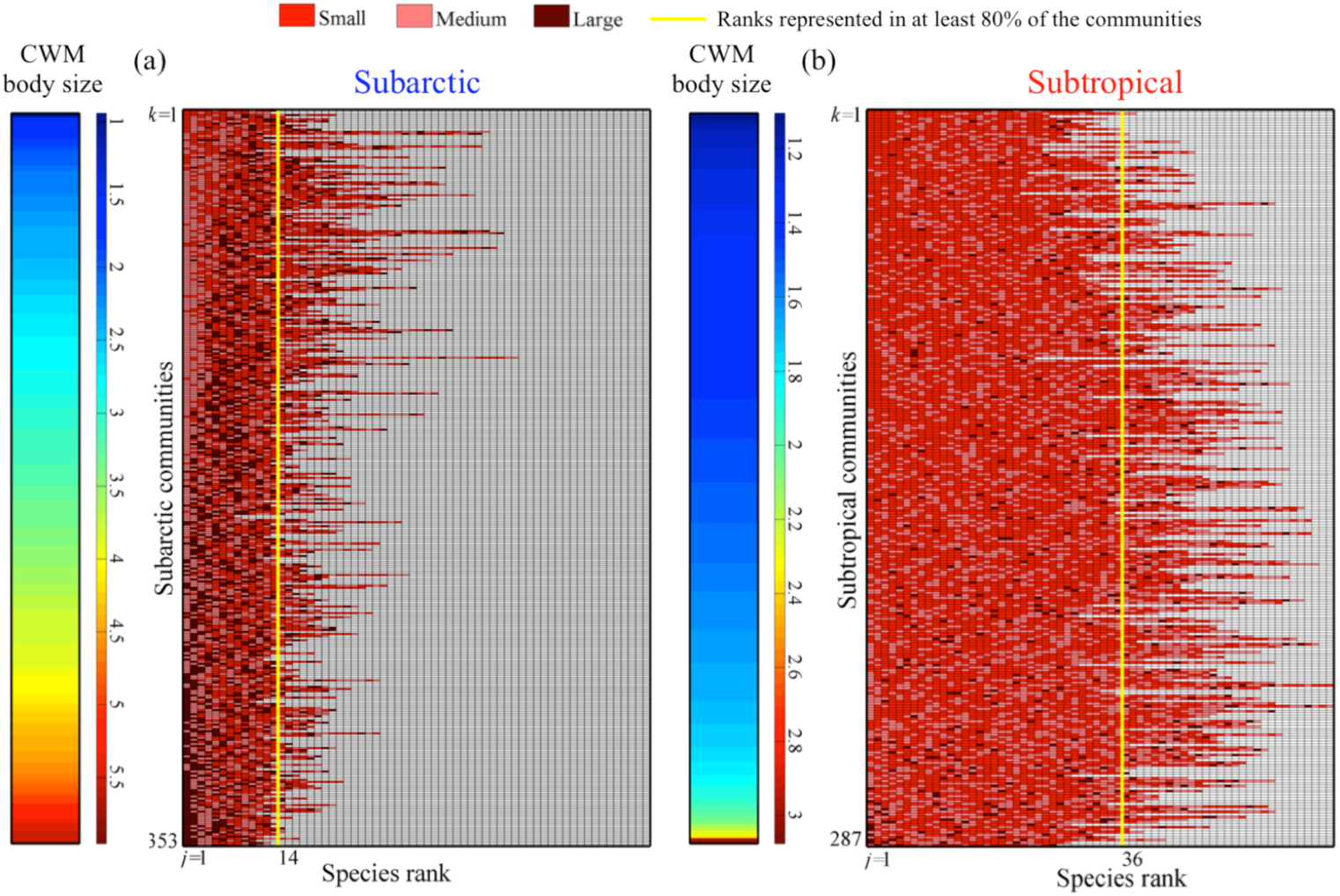
Species rank abundance distributions (RADs) with labelled body size categories of (a) subarctic and (b) subtropical communities ordered according to their community weighted mean (CWM) size (color bars).

Moreover, regarding rank 1 occupancy (the focus of our method), we observed that for similar values of CWM body size, the body size category occupying rank 1 varied (Fig. S7.2).

**Figure S7.2.**
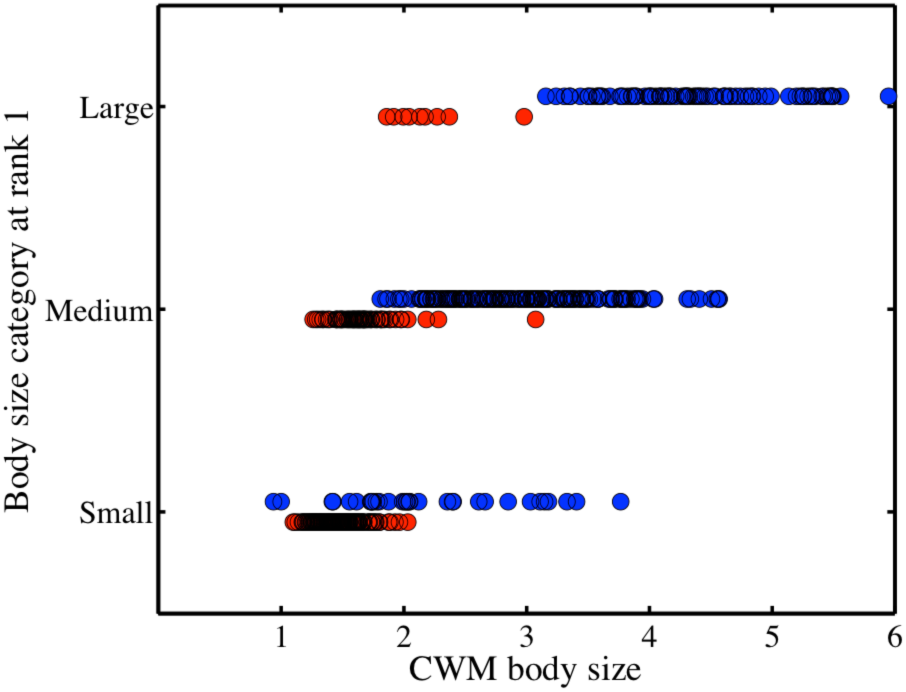
Relationship between community weighted mean (CWM) body size and body size category of the species occupying rank 1, for subarctic (blue circles) versus subtropical (red circles) communities.

In addition, we demonstrate the problem of using CWM body size to determine the dominant trait category. Specifically, we compared the dominant body size category determined according to the CWM body size (i.e. assigning one of the three body size categories of our study based on the CWM body size) with that determined according to the rank 1 in the RAD (i.e. the category currently dominating in the community). We then computed the error rate of miss-assignment of CWM approach; that is, for each of the three size categories in our study (i.e., total length < 1.38 ≥ 3.71 mm), the error rate corresponds to the number of communities in which the CWM body size corresponded to a category different from the category occupying rank 1 in that community. We then evaluated the error rates of CWM body size by comparing in a contingency table the size category of the CWM size body versus the size category occupying rank 1. For example, we found that, for the subarctic metacommunity, in 40% of the communities that the dominant category based on CWM body size was large, while the category occupying rank 1 was actually not large (Fig. S7.3). For the subtropical metacommunity, in 60% of the communities the dominant category based on CWM body size was medium, while the category occupying rank 1 was actually not medium (Fig. S7.3).

**Figure S7.3.**
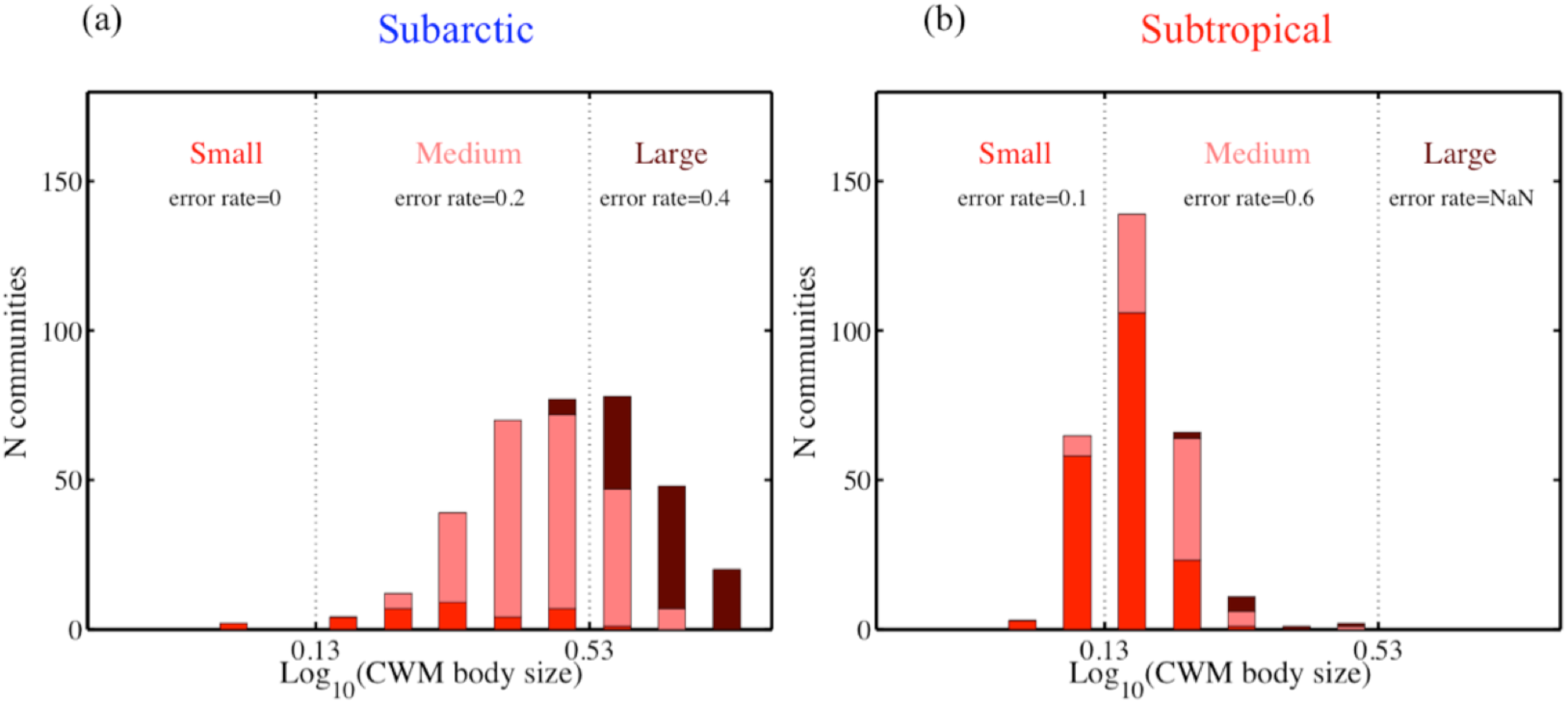
Distribution of rank 1 occupancy of the three body size categories in the size spectrum of log^10^(CWM body size). Error rates correspond to the rates of mismatch between the size categories determined by the CWM body size versus the size category occupying rank 1.

The overall error rate of the CWM body size was 0.26 (90/350) in the subarctic metacommunity, and 0.51 (146/287) in the subtropical metacommunity. The greater error rate in the subtropical metacommunity is due to the greater proportion of rare species in this region. Thus, we conclude that community-integrated measures are not suitable to test if a trait category affects species relative fitness and therefore dominancy.

## Appendix S8: Confidence intervals of *F-pool* via bootstrapping

In order to estimate the accuracy of the top-category *F-pool,* we applied bootstrapping (Efron & Gong 1983).

We bootstrapped 999 times for the top-category *F-pool* and estimated its 95% confidence intervals (C.I.). As such, 95% C.I. of the *F-rank_1_*/*F-pool* were also estimated. The C.I. of the top-category *F-pool* were narrow, and accordingly the relative frequencies of species with the top-category presented a unimodal distribution in both metacommunities.

**Figure S8.1.**
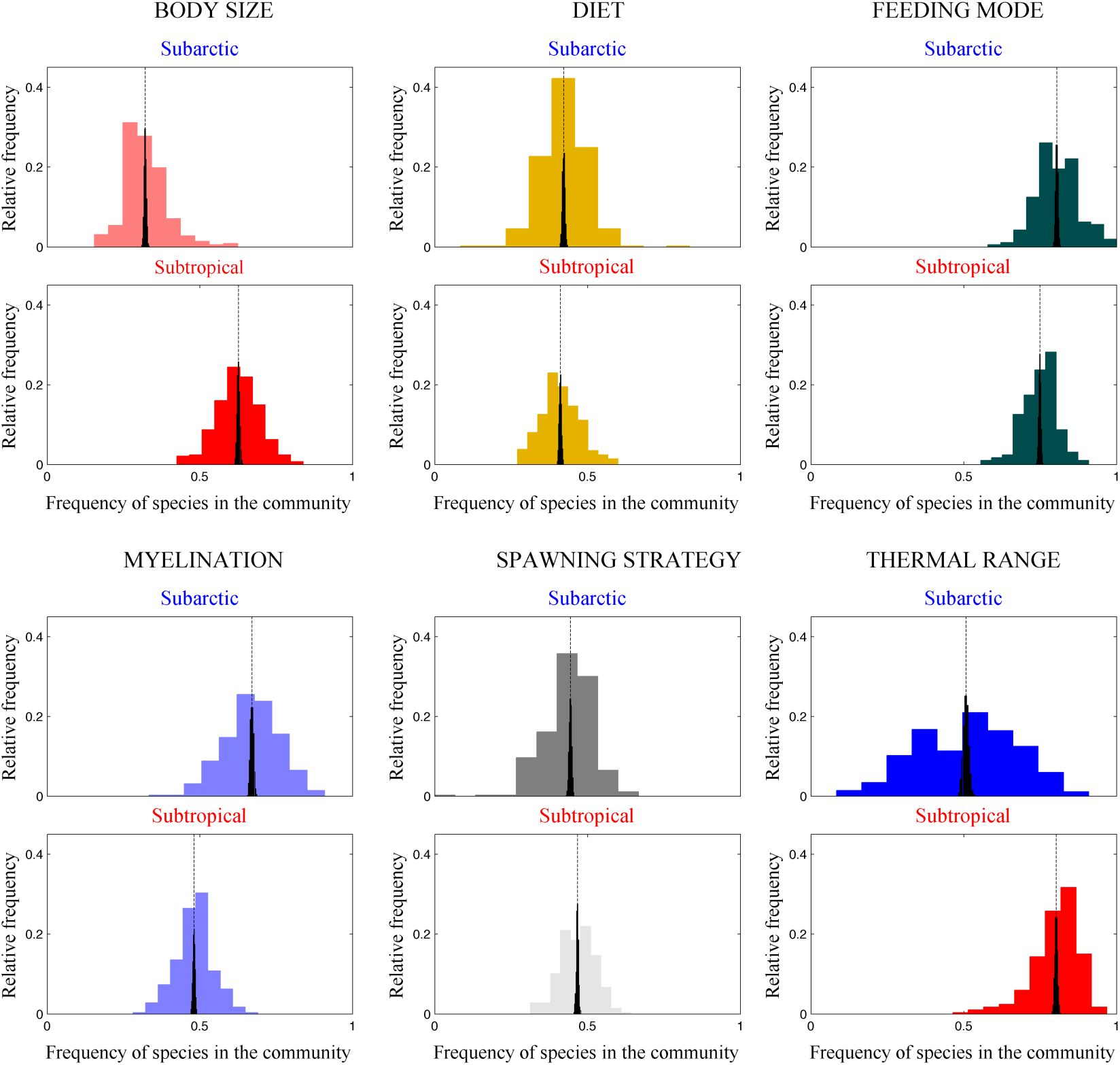
Histograms depicting the distribution of the relative frequency of species with the top-category in the set of communities (colour represents categories as in Fig. 2 and Fig.3 in the main text). The vertical dotted line corresponds to the estimated *F-pool* (i.e., mean of the distribution) and the black histogram shows the distribution of the 999 bootstrapped *F-pool*s used to calculate its 95% C.I.

